# Optimal timing for cancer screening and adaptive surveillance using mathematical modeling

**DOI:** 10.1101/2020.02.11.927475

**Authors:** Kit Curtius, Anup Dewanji, William D. Hazelton, Joel H. Rubenstein, E. Georg Luebeck

## Abstract

Cancer screening and early detection efforts have been partially successful in reducing incidence and mortality but many improvements are needed. Although current medical practice is mostly informed by epidemiological studies, the decisions for guidelines are ultimately made *ad hoc*. We propose that quantitative optimization of protocols can potentially increase screening success and reduce overdiagnosis. Mathematical modeling of the stochastic process of cancer evolution can be used to derive and to optimize the timing of clinical screens so that the probability is maximal that a patient is screened within a certain “window of opportunity” for intervention when early cancer development may be observable. Alternative to a strictly empirical approach, or microsimulations of a multitude of possible scenarios, biologically-based mechanistic modeling can be used for predicting when best to screen and begin adaptive surveillance. We introduce a methodology for optimizing screening, assessing potential risks, and quantifying associated costs to healthcare using multiscale models. As a case study in Barrett’s esophagus (BE), we applied our methods for a model of esophageal adenocarcinoma (EAC) that was previously calibrated to US cancer registry data. We found optimal screening ages for patients with symptomatic gastroesophageal reflux disease to be older (58 for men, 64 for women) than what is currently recommended (age > 50 years). These ages are in a cost-effective range to start screening and were independently validated by data used in current guidelines. Our framework captures critical aspects of cancer evolution within BE patients for a more personalized screening design.

**Significance:** Our study demonstrates how mathematical modeling of cancer evolution can be used to optimize screening regimes. Surveillance regimes could also be improved if they were based on these models.

**Graphical Abstract:** 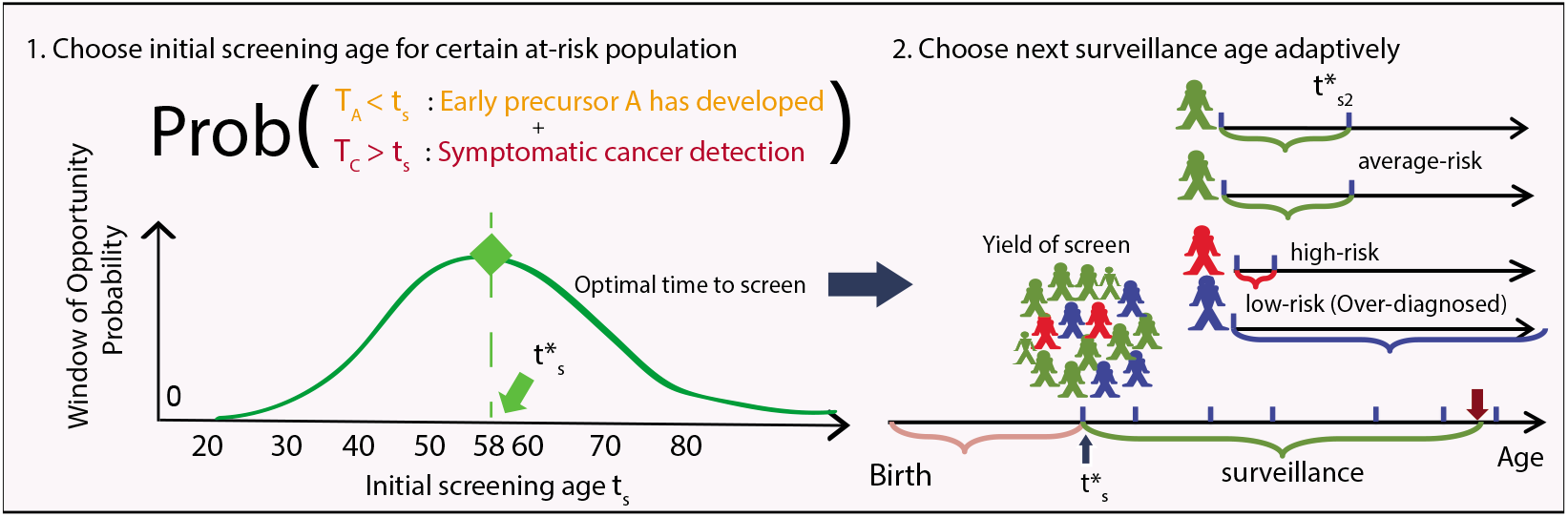

## Quick Guide to Equations

Ideally, a sensitive clinical screen will be offered to a patient during an opportunistic age window such that some event *A* (e.g., premalignant disease onset) has likely already occurred but an event *B* (e.g., cancer detection) has not yet happened. The rates of these events can depend on various risk factors (e.g., sex, ethnicity, environmental exposures) but the derivation of timing is universal. To maximize the probability that a patient is between *T_A_* and *T_B_* years of age at time of screening/surveillance *t_s_* is equivalent to simultaneously (1) maximizing Pr[*T_A_* ≤ *t_s_*] to ensure that event *A* has already occurred while (2) minimizing Pr[*T_B_* ≤ *t_s_,T_A_* ≤ *t_s_*] so that screens are not recommended when it is too late for an early intervention. This idea is reflected mathematically in the following relationships, which we will use in the optimal screen design methodology for quantifying ‘screening success’.

- Optimal screening age for some weight *w* on an adverse outcome, before time of patient death 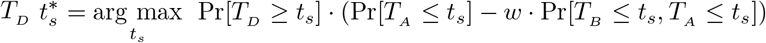
- Associated risk of adverse outcome for screening at time *t_s_ R_B_* (*t_s_*) = Pr[*T_B_* ≤ *t_s_*]
- Overdiagnosis rate - detecting precursor that will not become cancer by end of follow-up (age) *T_f_ OD*(*t_s_*) = Pr[*T_A_* ≤ *t_s_*|*T_B_* > *T_f_*]
- Successful diagnosis rate (i.e., early detection of future case of *B*) by time (age) *T_f_* > *t_s_ SD*(*t_s_*) = Pr[*T_A_* ≤ *t_s_*|*t_s_* < *T_B_* < *T_f_*]
- Over-screening - proportion of overdiagnosed cases in yield of *A* at screening time 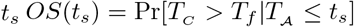
- Cost function *C*(*t_s_,T_f_*) associated with screening time *t_s_* and surveillance until a final age *T_f_ C* (*t_s_,T_f_*) = *OS* (*t_s_*) · (*T_f_* – *t_s_*)
- Associated risk for adaptive follow-up interval, conditional on screening/surveillance result (SR) at time *t_s_i__ R_B|SR_*(*t*_*s*_*i*_+1_) = Pr[*T_B_* ≤ *t*_*s*_*i*+1__|SR(*t_s_i__*)]

## Introduction

The main rationale for cancer screening is that earlier detection of a disease during a patient’s lifetime offers the opportunity to change its prognosis. Compared with incidental cancers, screening can improve the prognosis for patients with early occult cancers detected before symptoms develop, and perhaps even more importantly through removal of pre-invasive lesions such as adenomas in the colon, cervical intraepithelial neoplasia (CIN) in the cervix, and ductal carcinoma in situ (DCIS) in the breast. In 2018, it was estimated that nearly half of all cancer incidence in the US was attributable to detection by screening [1] but it is difficult to assess how many of these patients were destined to die from other causes before these cancers would have led to a symptomatic diagnosis, thus proving their screens to be ineffective. Current programs suffer from the problem of overdiagnosis of benign lesions and yet, with often dismal and costly consequences, there is still under-diagnosis of dangerous lesions because they were either missed by the screen itself or there was a lack of screening uptake by high risk individuals at the appropriate age [2]. Consequently, improvement in screening success is an important health policy research area, and one primed for quantitative assessment. Simply put, there is a *mathematical balancing act* to solve for human populations from both public health and cost-effectiveness perspectives: the goal is to maximize successful prevention of future lethal cancers (often by removing their precursors detected on a screen) and minimize the likelihood of overdiagnosis.

From a biological perspective, malignant cells develop in the body originally from normal cells at birth through the evolutionary process of carcinogenesis [3]. This process of cancer evolution is stochastic, meaning that random mutations can occur in cells throughout life that may eventually lead to a malignant phenotype selected in a tissue, but not necessarily so. Mathematical models of carcinogenesis, tumour progression, and metastasis are powerful tools to describe this process and, when applied to data, can infer important parameters for evolution such as time of metastatic seeding and selection strength of certain mutations that may confer resistance to treatment [4, 5]. Because such variables are not immediately measurable from data, mathematical models are now being explicitly incorporated into translational aims such as interval planning for adaptive therapy trials in cancer patients [6, 7]. We propose that these quantitative models can be further utilized at an earlier stage for cancer prevention - namely, to infer sojourn (waiting) times for specific stages of cancer evolution that we wish to target, and then to use this information to optimize the efficacy of cancer screening strategies and adaptive surveillance for early detection [8].

Clinical detection of biomarkers on a screen that alert us to phenotypic changes along the path to cancer (e.g., premalignant adenomas found during colorectal cancer (CRC) screening) are a snapshot of *field cancerization,* wherein groups of cells have acquired some but not all of the phenotypes necessary for clinical malignancy [9]. In CRC, the single first malignant cell creates a “Big bang” of tumor initiation, wherein the new malignant clone that develops already has all of the selective advantage it needs to progress, and random mutations accumulated are essentially neutral from an evolutionary perspective [10]. Before this occurs however, we know that there are certain stages of colorectal carcinogenesis that are both common in *type* (*APC* gene inactivation for initiation of adenomas) and in *timing* (adenomas become detectable around age 45 in males and females) [11]. Thus, although no two paths of carcinogenesis are exactly the same in two patients, there are most probable timescales for this stochastic process to take place (as seen in age-specific incidence curves) while still allowing for less likely events to happen (e.g., very young CRC cases).

These stochastic features were captured in the first mathematical descriptions of cancer, which have been studied and applied extensively in combination with biostatistical methods for over 60 years [12, 13]. By analyzing the structure of multistage models, we can also formulate these probability distributions as certain “windows of opportunity” that are crucial to capture in cancer screening planning, such as defining the likelihood to detect premalignant clones and simultaneously the likelihood that malignant clones would be unlikely to have developed yet. In this study, we derive conclusions from the theory of stochastic processes applied to carcinogenesis to formally optimize the timing of initial cancer screens in a population, and optimize subsequent surveillance adaptively, conditional on a previous exam result.

Here we present our analytical results for a generalized framework and then apply it in a proof-of-concept study for current screening of Barrett’s eosphagus (BE), the metaplastic precursor to esophageal adenocarcinoma (EAC). Briefly, because BE itself is essentially asymptomatic, the majority of BE patients remain undiagnosed and thus most EAC cases are diagnosed at an advanced stage. This is unfortunate because, (1) mortality associated with EAC is very high (majority of diagnosed patients die within a year), and (2) prior diagnosis of BE is positively associated with improved EAC survival in patients [14]. Therefore US and UK gastroenterologists have been focused on positively identifying BE patients on screening endoscopy, and then recommending these patients to undergo lifelong endoscopic surveillance in order to detect any dysplasia or early cancer that may be removed, thus preventing future lethal EAC. However the clinical reality is that only 0.2-0.5% of people with BE develop EAC each year [15]. Thus, the majority of diagnosed BE patients will attain little benefit per follow-up exam and over-surveillance of BE patients with no evidence of premalignant high grade dysplasia (HGD) poses a costly problem due to lifelong surveillance recommendations (as frequently as every 3 years by current guidelines). Beyond cost-effectiveness for a small number of microsimulated screening ages [16], the *optimal* starting age for BE screening in at-risk subpopulations is unknown.

Some screening recommendations have been justified by microsimulation models [11] but most screening guidelines consider current epidemiological data alone and are ultimately proposed by and decided by experts. The clinical consequences of these decisions are recorded as population outcomes, and then further research determines how effective these recommendations proved to be. When appropriate randomized control trial data is lacking, as is commonly the case in cancer screening [17], we propose that the initial choice of screening design could first be determined using an optimization algorithm with a machine learning and/or model-based approach. To address this problem, we derive probability equations to be used for optimal screening/surveillance timing and risk estimation, and present results for BE screening ages using a calibrated mathematical model that reproduces US cancer incidence data. Our predictions are in a cost-effective range and were independently validated with endoscopic screening data used for current BE guideline rationale.

## Materials and Methods

Below we derive the mathematical framework to determine the optimal timing of screening and surveillance of ‘at-risk’ patients using stochastic models of carcinogenesis. Herein, *cancer screening* refers to initial testing for the presence of a specified premalignant/malignant change, and subsequent tests that are offered after a postive screen diagnosis are defined as *surveillance.* If the age at completion of lead time (i.e., the length of time between the early detection of a preclinical state and the future clinical, symptomatic detection) surpasses patient lifetime, this patient will be considered an overdiagnosed case for that cancer. Published estimates of clinical endpoints like cancer incidence and the prevalence of precursors are reported on a population level, thus providing measures for mean dwell times in certain preclinical disease states that can be measured from population screening. For cancers with detectable precursors, we seek the age for which the probability is maximal that 1) individuals in a population harbor a screen-detectable premalignant clone, and 2) this timing occurs before incidental cancers have developed (Figure 1A).

**Figure 1:**
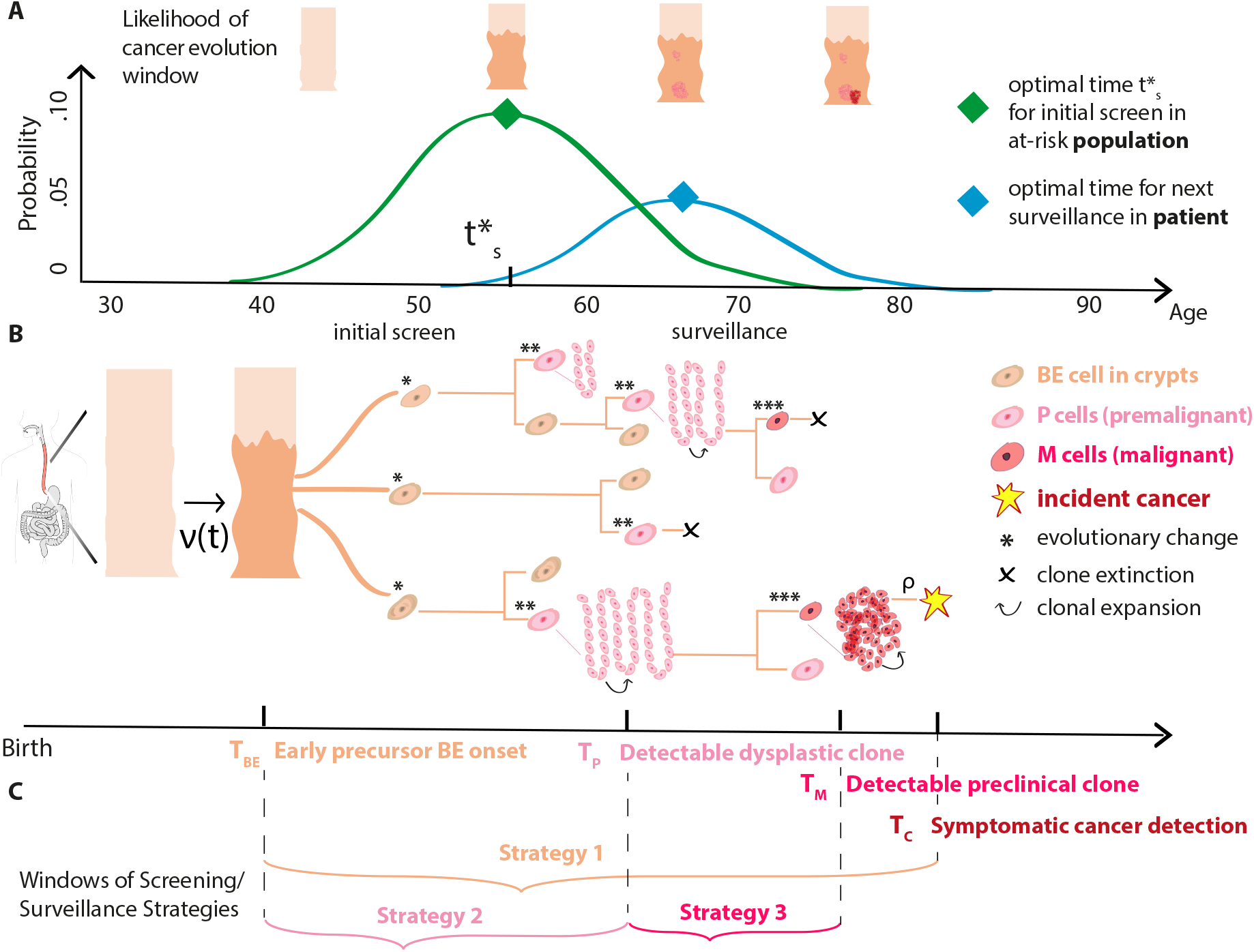
Optimizing screening and surveillance using branching process models of cancer evolution. A) The goal of early detection and prevention is to perform an initial screen on individuals within a larger population after premalignant changes reside in cells but before incidental cancer occurs in order to intervene. After an initial screen at time 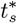, adaptive recommendations for patient-specific follow-up surveillance can iteratively account for the heterogeneity in a screened population. B) Somatic evolution of stem cell lineages since birth leads to stochastic trajectories (certain symmetric division examples shown on tree nodes) that accumulate mutations and may be selected for advantageous phenotypes. For example, within an at-risk patient’s esophagus, normal squamous epithelium may transform to a columnar BE segment with rate *ν*(*t*). Stochastic appearance of premalignant *P* cells can occur (e.g., after ‘two-hit’ gene inactivation of *TP53* in a cancer-promoting miroenvironment), which can then undergo clonal expansion described by birth-death-mutation processes. Malignant cells *M* that are initiated, in turn, may undergo clonal expansions or go extinct. Clinical detection of *M* clones that do not go extinct may occur through a size-based detection process with rate *ρ*. C) Criteria for optimal screening strategies can be formally defined using probability distributions in (A) for the timescales that characterize certain stages of carcinogenesis (see Text for details).

Biological processes not revealed in population data alone give rise to considerable inter-patient heterogeneity during the progression from normal tissue to incident cancer, and thus optimal timing for follow-up is expected to be highly heterogeneous within the population based on individual screening outcome. To account for this heterogeneity along with the temporal effects of screening (e.g., different prognoses resulting from screening earlier versus later), mathematical models can employ a stochastic approach to incorporate premalignant and malignant clonal expansions during clonal evolution explicitly (Figure 1B) [18, 19, 20, 21, 22]. This offers the advantage that the onset of (detectable) precursor clones can be defined as random variables and clonal population sizes can be tracked over time so that dwell times, risk of future cancer, and adaptive surveillance intervals may be formally derived (Figure 1C).

We first use basic tenets of probability theory to quantify these timescales in equations that can be applied for cancers that have measurable precursors to target (such as adenomas in the colon). The methods are organized in four parts: (1) General equations for optimal screening and application to multistage models of cancer, (2) Three strategies for optimal screening, (3) Metrics for efficacy assessment, and (4) Adaptive surveillance for 2 potential (negative/positive) screening outcomes. Then we apply the framework to determine the optimal timing of BE screening and surveillance in Results.

### Optimal screening framework

#### Optimal design options for weighting adverse outcomes

For some event *A* and later event *B*, maximizing the probability that a patient is between ages *T_A_* and *T_B_* at time of screening *t_s_* is equivalent to maximizing Pr[*T_A_* ≤ *t_s_*] to ensure that event *A* has already occurred while simultaneously minimizing Pr[*T_B_* ≤ *t_s_, T_A_* ≤ *t_s_*] to ensure that event *B* has not yet occurred by time *t_s_*. This statement is reflected mathematically as,

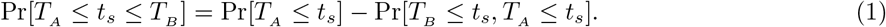

If event *B* implies event *A*, the second term on the right hand side of this equation simplifies to Pr[*T_B_* ≤ *t_s_,T_A_* ≤ *t_s_*] = Pr[*T_B_ ≤ t_s_*], the absolute risk of event *B* by time *t_s_*.

It may not be a justifiable public health goal to equally weigh the criteria that both event *A* has occurred and event *B* has not occurred, as is the case for maximizing the probability in Eq. (1). In general for events *A* and *B*, we define the probability of a “successful screen” with some specified positive weight *w*,

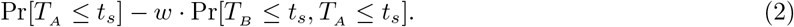

When *w* = 1, as in Eq. (1), both events contribute equally during optimization that a screen time occurs within the specific disease state window. As an alternative to equal weighting, Dewanji et al. [23] considered a simple illness-death model and maximized the probability that time *t_s_* be in the window between events *A* and *B* (where event *B* implies event *A*) with an extra penalty term for event *B* (which in our notation corresponds to the case *w* = 2).

If *w* > 1, maximization of Eq. (2) will yield earlier optimal screening ages 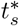 because there will be greater weight to avoid event *B* before screening, while older screening ages will be selected as optimal for *w* < 1. Cancer screening policy makers must decide how much weight should be given for each potential screening age *t_s_*, i.e., how heavily to guard against events *B* in the general population, such as cancer, balanced against the prospect of futile screens when patients have not yet undergone event *A*, such as premalignant onset.

#### Incorporating realistic time constraints for screening

Next we incorporate realistic cutoff times due to life expectancy to formally condition that we do not optimize screening ages that are unrealistically old, i.e., when the patient is more probable to have already died than to have developed the disease we wish to detect early enough to intervene. Here we include life table estimates for all cause mortality survival probabilities in the US [24] as an independent random variable *T_D_* in order to calculate Pr[*T_D_* ≥ *t_s_*], which is a close approximation when cause-specific cancer death rates in the general population are low, as is the case for esophageal adenocarcinoma. Thus, this method allows for random truncation due to death from other causes. For the general weighting *w* > 0 for events *A* and *B*, we compute the criteria defined as,

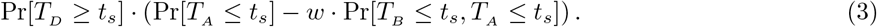

In these methods we will derive optimal screen times 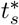 by maximizing Eq. (3) and refer to these 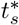 as the screening times that satisfy the *optimality criterion* for a given value of weighting factor *w*,

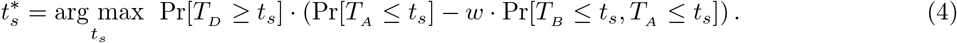

#### Quantifying risk for screening times

To perform a constrained optimization problem with additional risks incorporated, we quantify a penalty for certain screening times by introducing an equation for the risk 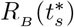 that some later event *B* has already occurred given some optimal screening time 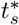,

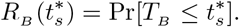

For cancer screening guideline decision-making, we can increase the weight *w* in Eq. (3) to find optimal time 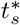 such that 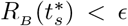, where *ϵ* is a threshold parameter for the allowable amount of risk of event *B*. Alternatively, if the risk of possibly allowing an undesirable event *B* before a proposed time 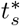 is too small to be of considerable worry, we reduce the weight *w* assigned to event *B* and thus raise the risk of event *B* to an appropriate, or tolerable, level.

#### Applying design to multistage carcinogenesis models

We derive optimal screening scenarios in a case study of Barrett’s esophagus (BE) using the multistage clonal expansion for esophageal adenocarcinoma (MSCE-EAC) model that has been employed within the Cancer Intervention and Surveillance Modeling Network (CISNET), a National Cancer Institute consortium, for cost-effectiveness studies of EAC prevention (see [25, 26, 27, 28] and Supplementary Figure S1 for model details). The random variables described in the next section will refer to stages of progression to cancer appropriate for BE but could be analogously formulated and expanded for similar multistage models of screening in colon, lung, breast, and others.

#### Random variables for screening outcomes

The random variables of interest that represent four clinical points of the multistage process are:

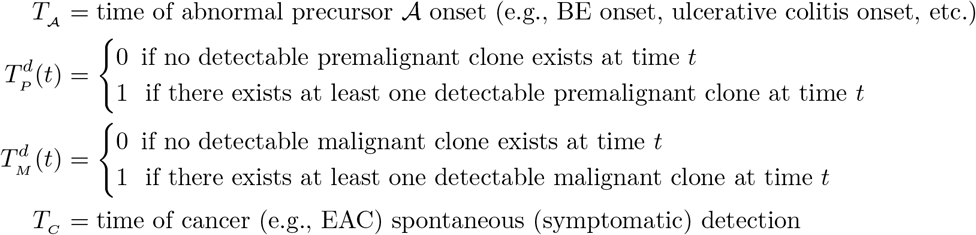

As applied in [25, 27], we derive the forthcoming formulas using a general precursor 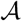 onset distribution 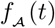,

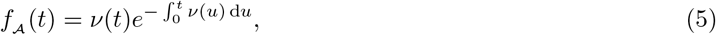

where *ν*(*t*) is the rate of precursor development. As has also been applied recently in case studies of gastric, lung, and oral cancers [29], we allow *ν*(*t*) to be time-dependent to allow straightforward incorporation of etiological agent prevalence that affect initiation rates in a population. For BE, we model the density *f_BE_* (*t*) with rate *ν*(*t*) given as a function of the prevalence of symptomatic gastroesophageal reflux disease (GERD) at age *t*,

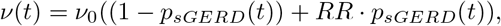

where *P_sGERD_* (*t*) is the prevalence of GERD symptoms at age *t, RR* is the relative risk of BE initiation due to exposure to GERD symptoms, and *v*_0_ is a positive constant.

Unlike the one-time occurence for 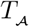 and *T_C_*, the detectability of premalignancy 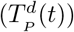 and malignancy 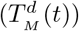 require specific definitions for appropriate binary random variables. If we designate these times to be the time of initiation or malignant transformation of a cell regardless of its fate (i.e., the ancestor to a clonal progeny that eventually goes extinct or that survives), we may derive analytical probabilities with first-passage time for continuous random variables *T_P_* and *T_M_*, respectively, using master equations. However, it is more clinically relevant to define these random variables as binary outcomes for detectable clones at the time of screening *t_s_*. In the derivations of our screening windows, the probability of detection at time *t_s_* of a premalignant clone that began with a single ancestor *P* cell initiated at time *s* is denoted as 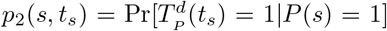, and analogously the probability of detection at time *t_s_* of a malignant clone that began with a single ancestor *M* cell initiated at time *s*, 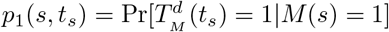.

In the Results below, we assign these binary detection variables the value 1 if a clone born at time s is non-extinct at time *t_s_* (size of clone is ≥ 1) and 0 otherwise. In the case of EAC, this “perfect sensitivity” definition will be useful as progressively better high-resolution imaging technologies and minimally invasive sampling devices are tested in trials in BE [16, 30] and used in future clinical practice. Lastly, we present the two survival functions that will appear throughout the mathematical derivations: 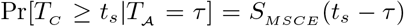, and Pr[*T_C_* ≥ *t_s_*] = *S_C_* (*t_s_*), where *S_MSCE_* (*t*) is the survival function for the multistage clonal expansion model of interest after precursor 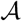 onset and *S_C_*(*t*) is the survival function for the full model (*S_EAC_* (*t*) for the MSCE-EAC model, see Supplementary Methods 1).

### Strategies for optimizing initial screening ages

#### Strategy 1: Optimization of precancerous yield before cancer

This strategy for initial screening success aims to maximize the probability that an individual has developed the precursor of interest before the time of screening but has not yet developed clinical cancer (Figure 1C, beige),

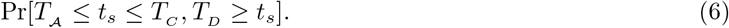

Thus to satisfy the optimality criterion in this scenario for weighting parameter *w* described by Eq. (3), we solve for the optimal screening age 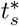 (Eq. (4)) as follows

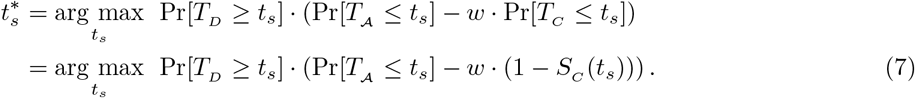

#### Strategy 2: Optimization of precancerous yield before dysplasia

This strategy for initial screening success aims to maximize the probability that an individual has developed the precursor before the time of screening but has not yet developed a detectable premalignant clone (e.g., dysplasia, Figure 1C, light pink),

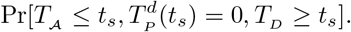

Then solving the optimality criterion for weighting parameter *w* on dysplasia development, we solve for this optimal screening age 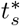 as follows,

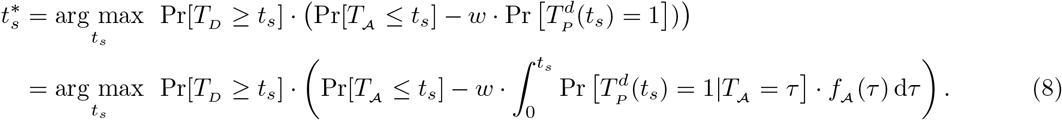

We utilize the filtered Poisson process (FPP) approach to analytically solve for 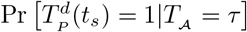 in the integrand (see Supplementary Methods 2 for derivation). Thus together with 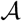 onset distribution in Eq. (5) and mortality risk, we have the analytical solution for optimal screening age in Eq. (8).

#### Strategy 3: Optimization of dysplasia yield before cancer

As a final example, we aim to maximize the probability that an individual has developed detectable dysplasia before the time of screening but has not yet developed a synchronous screen-detectable malignancy (Figure 1C, dark pink),

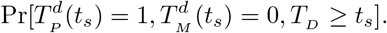

Thus the optimal screening time 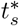 for Strategy 3 for specified *w* is given by,

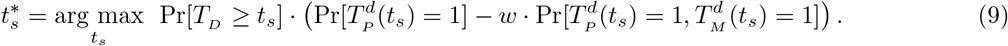

Integrating over all possible onset times for precursor 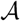, we can also derive the analytical solution for this optimal screen time (full derivation in Supplementary Methods 3).

### Metrics for assessing screening efficacy

#### Successful screening of future cancers cases

For screening effectiveness results, we define a successful diagnosis function *SD*(*t_s_*) for the probability of successful screening for future cancers before an age *T_f_*,

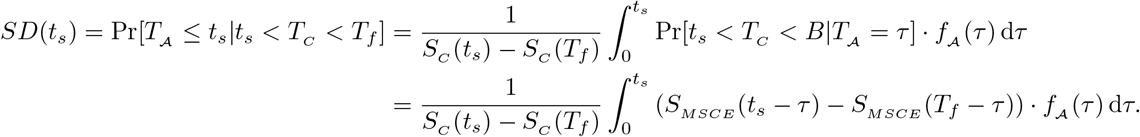

Thus *SD*(*t_s_*) is the proportion of patients who get cancer before *T_f_*, who biologically have 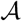 (undiagnosed) before screening age *t_s_*. Here we distinguish *T_D_*, a random variable for patient-specific death, from *T_f_*, a known constant time point (e.g., age 80) representing a set length of patient follow-up for comparison.

#### Overdiagnosis at screening age

Alternatively, *OD*(*t_s_*) is the probability of overdiagnosis (OD), given by

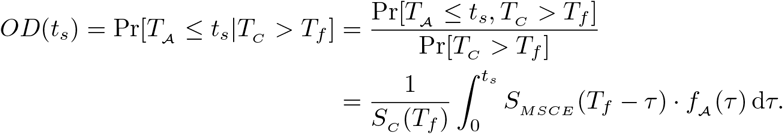

Thus *OD*(*t_s_*) is the proportion of patients who will not get cancer before age *T_f_* but who have 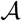 at time *t_s_*.

#### Proportion of over-screening in a positive-screen population

Another important quantity is the proportion of positive screens at the time *t_s_* who will undergo needless, costly surveillance. We can reformulate this proportion as a function with terms previously computed from *OD*(*t_s_*) above and Strategy 1 (Eq. (6)),

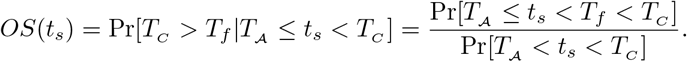

#### Proportion of successful screening in a positive-screen population

Lastly, the fraction of those who will be a successful screen (SS), for whom surveillance may be life-saving due to early detection of small cancers that will form prior to symptomatic detection, can be considered the positive predictive value (PPV) of the screen. We can reformulate this proportion as the complementary conditional event of *OS*(*t_s_*) above,

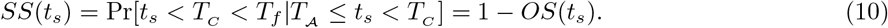

#### Comparing costs of different screening ages

To quantify the effects of different screening ages, we define a cost function *C*(*t_s_, T_f_*) for screening time *t_s_* and surveillance until an age *T_f_*. We aim to minimize the cost function which incorporates needless surveillance of the over-screened population from age *t_s_* to age *T_f_*,

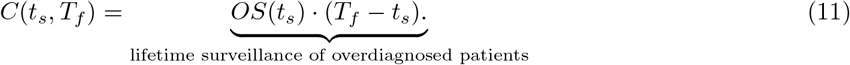

Here we assume that over-treatment/over-surveillance costs are likely proportional to the fraction of over-screened individuals multiplied by a set number of life-years after diagnosis. Assuming that within each strata there is equal risk of premalignancy to treat during surveillance, this weighting determined by number of surveillance years is reasonable in our cost metric.

### Optimizing re-screening and surveillance in an adaptive way

After a screen, current clinical guidelines recommend fixed follow-up examination intervals based on a crude analysis of existing evidence, often without incorporating known risk factors of patient progression (e.g., patient age, gender). In our proposed method for adaptive surveillance, mechanistic modeling of the carcinogenesis process is assessed *conditional* on patient demographic/clinical features and detected stage of progression (or lack thereof) at screening time *t*_*s*_1__ to provide a more refined surveillance recommendation.

#### Outcome 1 (negative for precursor): when to re-screen

After a normal screen, an open question is whether the patient should be asked to return for repeat screen at a later time. Analogous to Strategy 1, we could aim to screen this individual again at a later age *t*_*s*_2__ when he/she is most probable to have developed the precursor by this time but not yet clinical cancer,

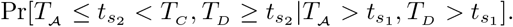

We derive the optimal age (see Supplementary Methods 4 for full solution) for a next screening time as,

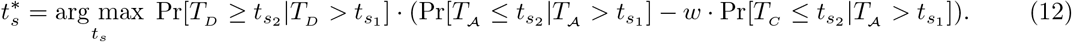

#### Outcome 2 (positive for precursor): associated cancer risk

After diagnosis, we can iteratively condition on the screening/surveillance result at time *t_s_i__, SR*(*t_s_i__*), and derive an optimal interval for the next surveillance given some chosen outcome of interest to obtain 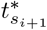. For Outcome 2 with no dysplasia/neoplasia detected, the subsequent risk for cancer developing before a suggested next surveillance exam at time *t*_*s*_2__ is,

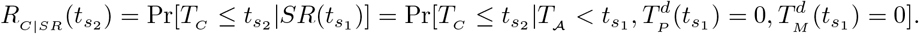

In general, a physician will often have no way of knowing how long a patient has harbored undetected cancer precursors in the body, only that onset occurred sometime before screening 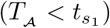. But if we could measure this precursor onset time *τ*, then his/her associated cancer risk for next surveillance age *t*_*s*_2__ would be more specific than defined above and would be defined (see Supplementary Methods 4 for derivation) as,

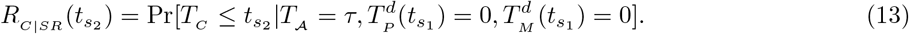

#### Outcome 2 (positive for precursor): Strategy 1

In ‘adaptive’ surveillance, we consider when the ideal next surveillance exam should happen conditional on the outcome of a screen or last surveillance exam. Assuming we have an estimate *τ* of a patient’s precursor onset time, we solve for optimal next-surveillance age 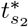 (see Supplementary Methods 5 for full solution),

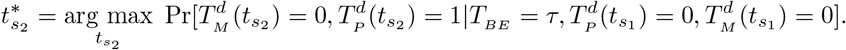

#### Previously calibrated parameter values reproduce population cancer incidence

The optimal screen design methodology presented thus far is applicable to multistage models, in particular branching process disease models with defined stages of clonal expansions occurring during carcinogenesis. In a case study for BE screening presented here, we apply the optimality criteria defined above for the MSCE-EAC model specifically.

For this model, the cell-level description is linked to the population scale by means of the model hazard function, defined as the instantaneous rate of detecting cancers among individuals who have not been previously diagnosed with cancer [27]. Briefly, the hazard function can be derived from the backward Kolmogorov equations for the stochastic multistage process described above and solved numerically via a system of coupled ordinary differential equations (Supplementary Methods 1). Thus, one may infer rates of cellular processes from cancer incidence data (see [25, 27] for model results using EAC incidence data from the Surveillance, Epidemiology, and End Results (SEER 9) registry 1975-2010, with predicted trends to 2030). For the quantitative Results in the use case below, we evaluate the equations defined in the above methods using values for evolutionary variables previously estimated in [27], provided in Supplementary Table S1.

## Results

Recommendations for risk-stratified Barrett’s esophagus (BE) screening and surveillance regimes have been proposed to optimize beneficial use of healthcare resources, but these have been limited in strata choices, variable across organizations, and based on conclusions from observational data alone [31]. For example, one current US guideline from the American College of Gastroenterology (ACG) strongly recommends BE screening for men with chronic and/or frequent (more than weekly) gastroesophegaeal reflux disease (GERD) symptoms and two or more risk factors for BE or EAC. These risk factors include: age greater than 50 years, Caucasian race, presence of central obesity, current or past history of smoking, and a confirmed family history of BE or EAC (in a first-degree relative) [32]. Alternatively in the UK, the British Society of Gastroenterology recommends endoscopic BE screening for any gender of patients with chronic GERD symptoms and multiple risk factors (at least three among: age 50 years or older, white race, male sex, obesity)[33]. After BE diagnosis, all of the guidelines recommend a certain uniform timing of surveillance and treatment regimes that are determined by presence or absence or detectable dysplasia (no other risk factors considered formally, including patient age), which were proposed by experts and deemed cost-effective [28].

Instead, we applied our framework in the case of BE to obtain optimal screening and surveillance design from theoretical predictions *a priori*. In our notation for this framework (Materials and Methods), we specifically refer to initial precursor development 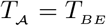, the detection of premalignant (high grade dysplasia) and preclinical malignant clones 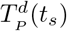 and 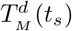, respectively, and clinical cancer occurrence *T_c_* = *T_EAC_*.

### Optimal ages to initialize screening in Barrett’s esophagus

We first used the model to optimize the choice of recommended age to start screening, rather than assuming age 50 as the threshold to begin screening across risk groups as guidelines suggest currently (in all those that specify an age [31]). We applied Strategy 1 (Materials and Methods) for a single screen to maximize the probability that an individual has developed BE before the time of screening but has not yet developed clinical EAC, thus obtaining optimal screening ages 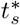 defined in Eq. (7) for weight parameters *w*.

In line with current risk factors, we computed optimality criteria for timing of individuals screened for BE stratified by age, sex, and GERD status (i.e., general population or symptomatic GERD population). Figure 2 depicts the contour plots of these optimality criteria for individuals born in year 1950 along with the optimal ages for each *w* (red lines). We sought ages by whole years for simplicity, in the way that guidelines are formulated currently. For *w* = 1 (i.e., equal weighting of positive screen and safeguarding from cancer before screening), the optimal screening times were (A) 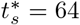 years old for males, (B) 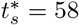 years old for males with GERD symptoms (indicated by red diamonds), (C) 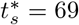 years old for females, and (D) 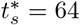 years old for females with GERD symptoms (indicated by red circles). In consistency with previous efficacy and cost-effectiveness analyses of BE screening [16, 25, 27, 28], we present results applied to the 1950 birth cohort as a ‘base case’ study and note that screening optimization predictions are robust through modern birth cohorts in a 50 year range (Supplementary Figure S2). This is consistent with previous BE prevalence estimates at index endoscopy that found only a small secular age-specific trend in index BE prevalence in white male GERD screenees but no trend in women across the calendar year period years 2000-2006 [34].

**Figure 2:**
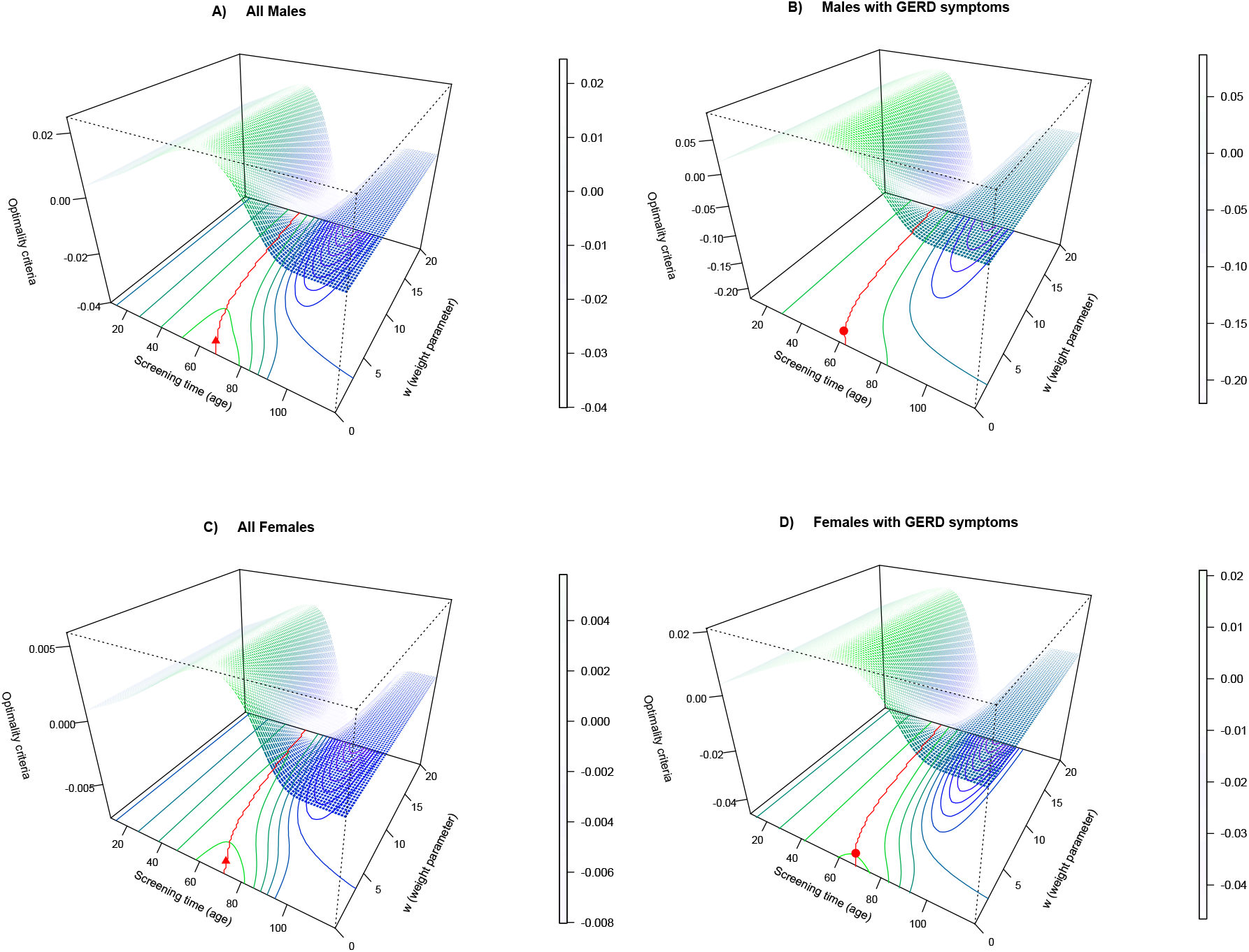
Contour plot of optimality criteria for single screen Strategy 1. The optimality criteria are plotted for given screening age *t_s_* between ages 10 and 119 and weighting parameters *w* ∈ [0, 20]. The red line on the 2D contour plot denotes the optimal screening ages for all races combined, born in 1950.

Table 1 provides the optimal ages computed for stratified populations along with associated metrics of efficacy. Optimal screening ages 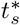 defined in Eq. (7) depend on a choice of parameter *w*, to protect against the adverse outcome of EAC development before age of screening. For males in the general population, the range of optimal screening times was 48-68 years of age (mean age 53 years) with *w* ∈ [0, 20] for Strategy 1.

**Table 1:**
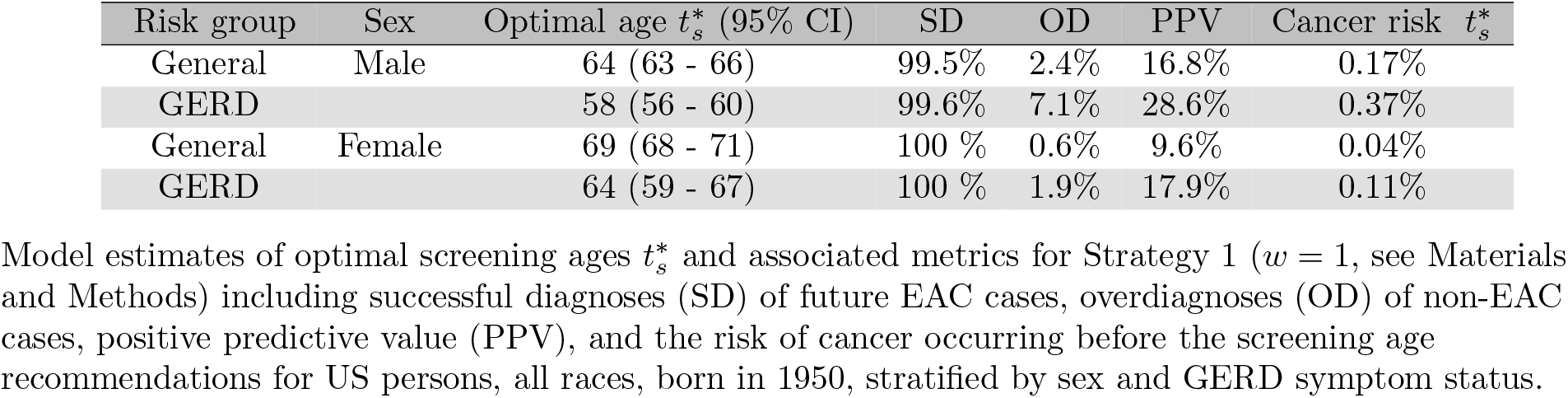
Optimal screening ages for BE in GERD and general populations

| Risk group | Sex | Optimal age $t_s^*$ (95% CI) | SD | OD | PPV | Cancer risk $t_s^*$ |
| --- | --- | --- | --- | --- | --- | --- |
| General | Male | 64 (63 - 66) | 99.5% | 2.4% | 16.8% | 0.17% |
| GERD |  | 58 (56 - 60) | 99.6% | 7.1% | 28.6% | 0.37% |
| General | Female | 69 (68 - 71) | 100 % | 0.6% | 9.6% | 0.04% |
| GERD |  | 64 (59 - 67) | 100 % | 1.9% | 17.9% | 0.11% |

Model estimates of optimal screening ages $t_s^*$ and associated metrics for Strategy 1 ( $w = 1$ , see Materials and Methods) including successful diagnoses (SD) of future EAC cases, overdiagnoses (OD) of non-EAC cases, positive predictive value (PPV), and the risk of cancer occurring before the screening age recommendations for US persons, all races, born in 1950, stratified by sex and GERD symptom status.

### Weighting of associated costs affects screening optimization

From a preventative viewpoint, screening recommendations may be aimed to reduce the risk that patients developed high grade dysplasia (HGD) prior to screening to avoid invasive radiofrequency ablative treatment after index endoscopy. To determine how optimal screening ages shift if we aim to screen patients who initially have BE but do not yet have existing HGD clones, we applied Strategy 2 in our optimization to obtain optimal screening times (see Eq. (8)).

Figure 3 depicts the contour plots of the optimality criteria for individuals born in 1950 stratified by age, sex, and GERD status along with the optimal ages for each *w* (red lines). For *w* = 1 (i.e., equal weighting of positive screen and safeguarding from HGD before screening), the optimal screening times were 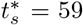 years old for males and 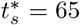 years old for females, which are 5 and 4 years earlier for males and females, respectively, than the optimal ages computed for the general population in Strategy 1. Interestingly, for the symptomatic GERD population, there is no optimal time obtained for *w* = 1 because it is unfeasible to screen early enough such that those with GERD and BE do not have any small HGD clones present. However, to lessen the cost of having small HGD clones at time of screening in our optimization, we set *w* = 0.95 and found the same optimal ages for GERD patients as were found using Strategy 1.

**Figure 3:**
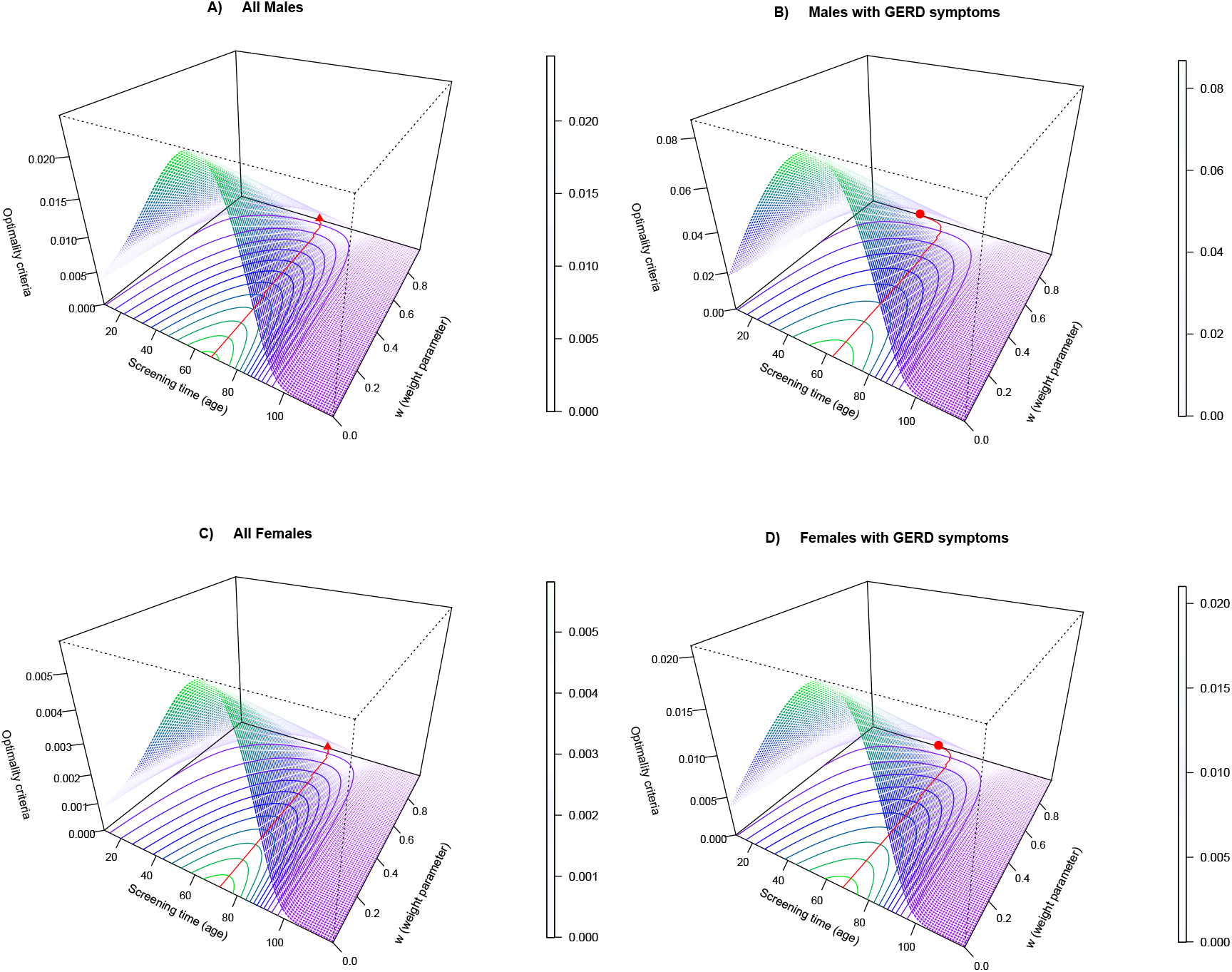
Contour plot of optimality criteria for single screen Strategy 2. The optimality criteria are plotted for given screening age *t_s_* between ages 10 and 119 and weighting parameters *w* ∈ [0,1). Note, as the weighting factor increases the optimal screen age becomes earlier. The red line on the 2D contour plot denotes the optimal screening ages for all races combined, born in 1950.

### Optimal screening ages can improve ad hoc guidelines by decreasing over-surveillance

We compared our optimal screening ages with current BE screening guidelines in the US and UK [32, 33] that recommend screening high risk male patients with GERD symptoms starting at age 50 and older. This choice of starting age was based on results from the US Clinical Outcomes Research Initiative (CORI study that quantified age-specific prevalence of BE in patients (mainly white race) with no prior EAC at time of first endoscopy, stratified by sex and presence of GERD symptoms as indicated in the endoscopy report [34] (Figure 4A).

**Figure 4:**
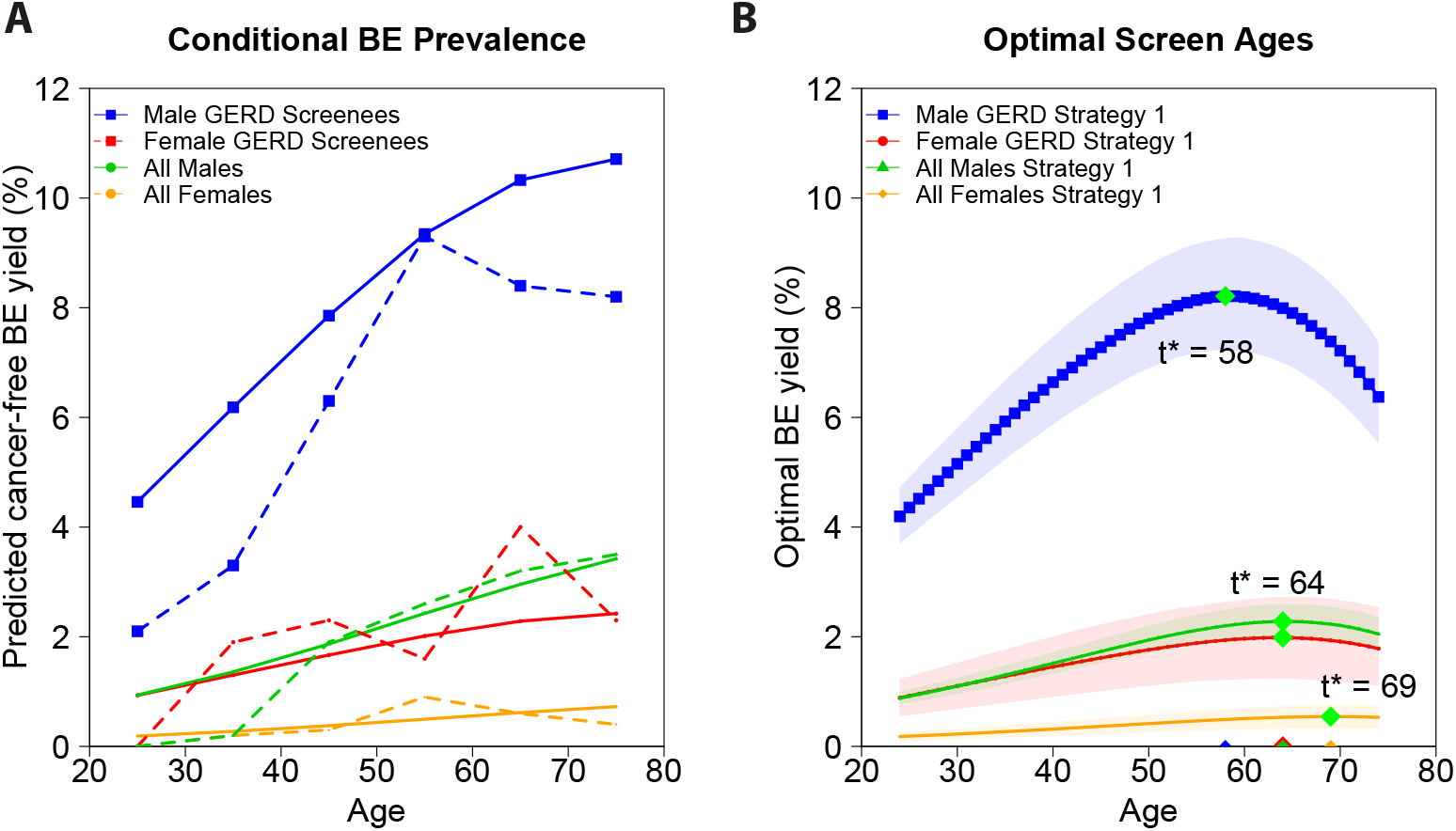
Validating predicted screening ages and computing optimal ages with Strategy 1. A) Model predictions of the conditional prevalence (solid lines, Eq. (14))) recapitulate CORI data for BE prevalence detected on index endoscopy in cancer-free individuals (dashed lines). B) Strategy 1 (single age results using Eq. (7) with *w* = 1, 95% confidence intervals shaded) incorporates both protection from previous cancer formation (missed cases) and all-cause mortality pre-screening time into optimal screening decision-making (green diamonds).

First we directly compared the CORI prevalence with our model by computing the analogous conditional probability to represent the fraction of first-time diagnosed BE cases by age within the total 155,641 cancer-free patient cohort who underwent first endoscopy between 2000-2006,

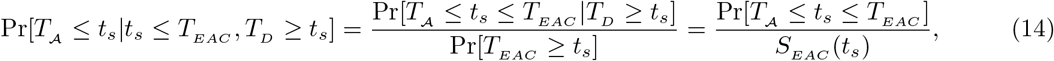

where this numerator was computed previously in Strategy 1 (see Eq. (6)).

This data independently validated the similar age trends predicted by our model (which was fit to SEER incidence, irrespective of CORI) for these strata born in 1950 (Figure 4A for *w* = 1). In only the case for the highest risk group, our estimates of prevalence were found to be slightly greater than the published estimates likely because the model calculates the true yield of all BE cases detectable by a highly sensitive screen rather than diagnosed BE cases among patients in the CORI study, which may have also suffered from a ‘harvesting’ effect for older age groups that contained less patients [34]. Further, we independently confirmed the two main Results from Rubenstein et al. that: (1) the yield of upper endoscopy for the diagnosis of BE rapidly increases among men with GERD until approximately age 55, and then the prevalence begins to level-off, and (2) women with GERD were at no increased risk compared with men without GERD (red and green lines, respectively) and thus, if screening were considered, it should be recommended at the same age (or not at all, if lacking additional risk factors). With the ability to compute not only this prevalence analytically but furthermore an optimal criterion with Strategy 1, we can maximize the BE yield at any continuous age *t* subject to a *joint* probability of being cancer-free and alive (Figure 4B for *w* = 1). For previous guideline formulations, such clinical decisions had to be made essentially ad hoc or *conditional* with cohort data grouped into decade age-groups [34] for proposals of most effective screening age for BE yield.

With our mathematical formulation, we can also quantitatively compare costs associated with chosen screening initialization ages. Using symptomatic GERD males as our example, weg compared the initial screen age of *t_s_* = 50 (current practice) versus our optimal age of 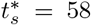. The costs associated with over-surveillance defined in Eq. (11) in detecting cancers up to age *T_f_* = 80 in these two cases was,

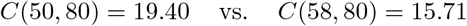

Thus, the overall costs associated with screening at optimal age 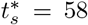 were estimated to be less than current costs, mainly due to savings gained from eliminating 8 years of futile surveillance in a large number of middle-aged patients. For both screening ages 50 and 58 in GERD men, there was less than 0.5% risk of screening too early (i.e., very few men who develop EAC will have developed BE after those screening ages and progressed undetected without surveillance). Also, screening younger ages did not always imply increased numbers of future cancer cases detected, the main goal for successful screening. The probability of a missed EAC case occurring in a male GERD patient between these two ages was minimal,

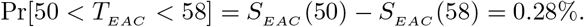

The current cost burden for EAC cancer control is then likely attributable moreso to years of futile surveillance than costs associated with missed cases expected to arise before later screening age recommendations.

### Less than 3% of normal patients will be found with BE on re-screen

For a first outcome, suppose a patient is screened for suspected BE and the screen at time *t*_*s*_1__ returns negative for BE (Outcome 1, Materials and Methods). An important decision for clinicians is whether to suggest that this patient come back for a one-time follow-up second screen at a later age and if so, at what age they should return. Current guidelines do not advise re-screening unless a patient is healing from esophagitis wounding to ensure underlying BE was not missed initially [32]. This recommendation was based on a CORI study that found 2.3% of patients with Outcome 1 had BE on repeat examination, implying that BE is rarely missed and develops early in patients on surveillance [35]. With our methods we can quantify: (1) when is the best age to offer a second screen, and (2) how likely it would be that BE would be found at that age.

We optimized when to screen this individual again at a later age *t*_*s*_2__ when he/she is most probable to have developed BE by this time but not yet clinical cancer. Given a range of prior screening times *t*_*s*_1__ ∈ {45, 50, 58, 64, 69}, Figure 5 shows results for optimal subsequent screening times *t*_*s*_2__, where *t*_*s*_1__ < *t*_*s*_2__ < 120. The black diamonds depict the optimal screen times 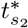 (see Eq. (12)) for this simulated cohort (US population, all races, born in 1950). For our choices of *t*_*s*_1__ in ascending order, we found that optimal follow-up screens were (A) 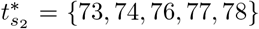 for all males combined, (B) 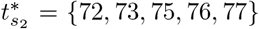 for GERD males, (C) 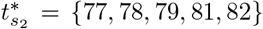 for all females combined, and (D) 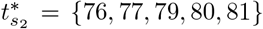 for GERD females. Thus, even though initial screening times may be over 20 years apart, the optimal range of a follow-up screen at age 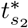 is only within a 5 year window.

**Figure 5:**
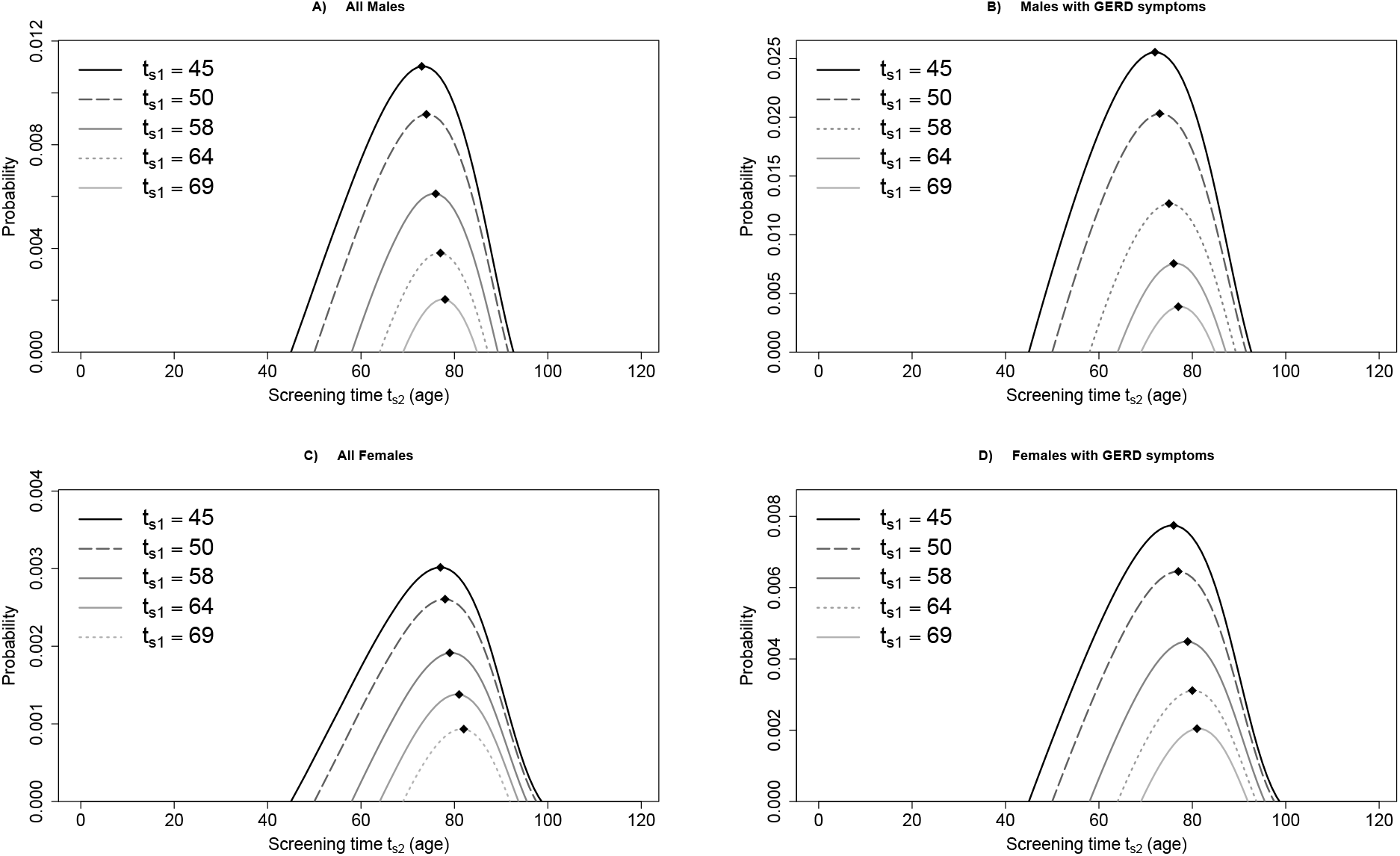
Optimality criteria for Outcome 1 (negative for BE): Strategy 1. The optimality criteria for each choice of next screening time *t*_*s*_2__, given prior screening age *t*_*s*_1__ ∈ {45,50,58,64,69} are denoted for weighting *w* = 1 (including typical case for current practice, *t*_*s*_1__ = 50, shown in dashed lines). Optimal re-screen times 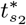, shown for all races, born in 1950 (diamonds), and for the risk group-specific optimal 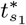 (dotted lines), were (A) 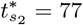 for all males, (B) 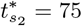 for GERD males, (C) 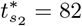 for all females, and (D) 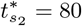 for GERD females.

Our prevalence figures for these are very close to those found in the CORI study for both genders [35], further validating our straightforward approach to assess public health questions that typically depend on grouped, conditional empirical population data (alive, cancer-free) for prevalences rather than optimally scheduled predictions of screening outcomes and yields based on continuous age. Importantly, we found that the initial age of screening (rather than time since initial screening) affects the future probability of BE yield; such information is not currently considered when clinicians decide whether certain GERD patients should return to be re-screened. For example we found that, if initial screening took place for GERD males at age 58, there would be a 1% chance of a positive re-screen rather than 2 – 3% when screening first at age 50. Importantly, even this 1% will be mostly overdiagnoses as *de novo* BE is much less likely to have time to develop to EAC within the remainder of patient lifespan, thus further devaluing a re-screen after initial negative result.

### Accurate cancer risk prediction depends on premalignant molecular age

In Outcome 2, a patient is found to have BE but no neoplasia (HGD/malignancy) at screening time *t*_*s*_1__. Current guidelines suggest these patients return for surveillance endoscopies every 3-5 years for the rest of their lives [32, 33] because they still carry risk of EAC that may in fact increase with time regardless of persisting without a diagnosis of dysplasia [36]. Furthermore, non-dysplastic BE patients with “missed” HGD/EAC were significantly older than those who progressed later [37], suggesting the important role age plays as a significant risk factor indicating missed EAC and the need for surveillance of BE patients who have unknowingly harbored BE for longer than younger patients. This prompts the question whether patient-specific BE “dwell time” influences future neoplastic risk and thus also the efficacy of timing of surveillance schedules.

If we measured BE onset time *τ* for a patient with Outcome 2 at time *t*_*s*_1__ (e.g., via methylation-based molecular clocks [26]), then his/her associated EAC risk for given next screening age *t*_*s*_2__, is given by Eq. (13). Figure 6 depicts EAC risk for US 1950 birth cohort, with prior screening ages *t*_*s*_1__ ∈ {45, 50, 58, 64} for males (A) and *t*_*s*_1__ ∈ {45, 50, 64, 69} for females (B). As shown in Figure 6, the model predicted that in general the associated EAC risk for those that developed BE earlier in life is greater than the risk for those BE patients who developed BE later in life. For the screen performed at time *t*_*s*_1__ = 50 (dashed lines) in both sexes, the BE patient who developed BE at age 20 (purple line) has a 2.4 predicted relative EAC risk versus the BE patient who developed BE at age 40 (yellow line). Currently in clinical practice however, these two patients would likely be treated equally because they were diagnosed at the same age. Our findings reiterate that heterogeneity in BE onset ages may translate to large differences in EAC risk [26].

**Figure 6:**
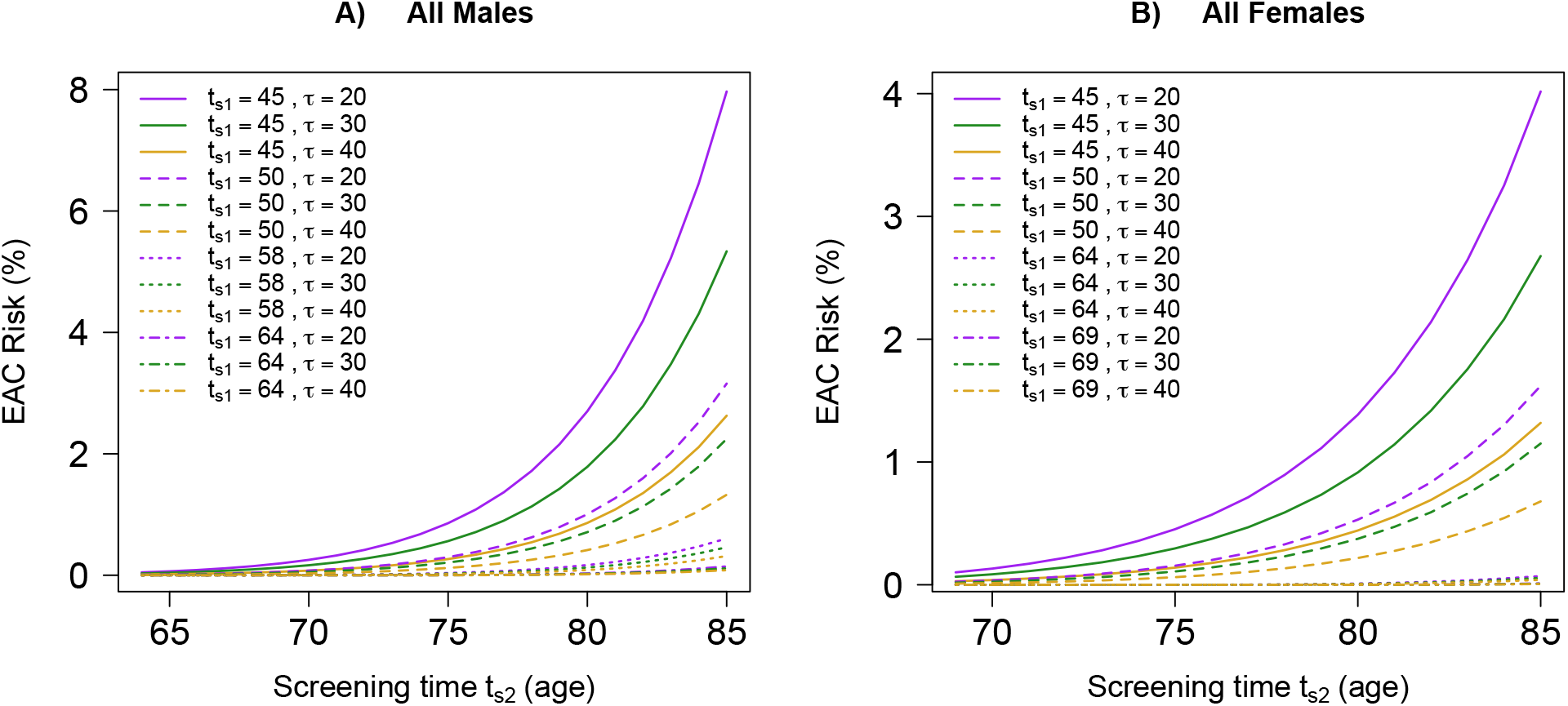
Associated EAC Risk for Outcome 2 (positive for BE). The EAC risk in Eq. (13) is plotted for each choice of next screening time *t*_*s*_2__ until age 85, given prior screening age *t*_*s*_1__ ∈ {45,50,58,64} (denoted by line type) for males (A) and *t*_*s*_1__ ∈ {45,50,64,69} for females (B), and a BE onset age *τ* ∈ {20,30,40} (denoted by color).

### Sensitivity of predictions

Our multistage model was calibrated to population incidence data [27] and has been previously tested to quantify the effects of perturbations in model parameters in Supplementary Table S1 such as initiation and growth rates of dysplastic clones that reflect published premalignant prevalence within BE (see details in [25, 38, 39]). Similarly, in these analyses, we tested our model for risk-stratified populations and found that our optimal timing predictions are robust to perturbations in parameters, as calculated for optimal ages in a bootstrap analysis of 1,000 re-sampling iterations of the Markov chain Monte Carlo (MCMC) posterior distributions for each calibrated parameter (Figure 4B, shaded regions).

We also considered 5 additional modern birth cohorts beyond the 1950 base case and found that: (1) Optimal screening ages remain unchanged for males (GERD and general populations) born between 1950 and 2000, and (2) similar results to the base case were found for females such that optimal screening times 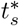 varied between these cohorts by only 3 years in females with GERD and 2 years for the general female population (Supplementary Figure S2). For optimal choice of adaptive surveillance based on screening outcomes however, we found that results will be sensitive to certain patient-specific parameters such as BE onset age and thus these will be important to include in order to perform more personalized surveillance.

### Data and code availability

All analytical distributions to recreate our Results and apply to other data are available in the Text and Supplementary Methods. Parameters used as input data for the case of BE screening are provided in Supplementary Table S1. Code to solve equations was developed in R (version 3.6.1). This framework will be made publicly available upon publication (github.com/yosoykit/OptimalTimingScreening).

## Discussion

Mathematical models of cancer evolution are being widely developed to understand timing of events and dynamics of treatment resistance. In this study, we applied such models to optimize screening and surveillance. Three examples of modeling techniques that could be formally assessed for screening timing include Markov models for natural history of disease [40, 41, 42], biologically-based models that incorporate dynamic processes like clonal expansions and biomarker shedding [18, 19, 22, 25, 43, 44], and biological event timing models that infer ordered genetic events [45]. While utilizing different data types, these all include distinct stages of latency periods separated by rate-limiting events on a patient’s forecasted trajectory. Our mathematical formalism provides one means of targeting these windows to answer early detection questions with models. Our approach also provides the flexibility of choosing the ‘window of opportunity’ and the specified weight *w* for the adverse event to suit the purpose of the health policy goal.

Several previous studies address optimal screening design via biomathematical modeling [18, 19, 23, 46], but this study targeted cancer precursors specifically and allowed for this flexible weighting of events to determine the specific age to screen (and whether to screen) in an analytical framework that does not require simulation. Microsimulation studies using models have also been used to inform policy-making decisions on clinical recommendations/modalities, for example in colorectal cancer [47] and lung cancer [40] screening, by the US Preventive Services Task Force (USPSTF). The model we present in our study has been previously used in this way to evaluate certain screening and surveillance scheduling of Barrett’s esophagues (BE) to prevent esophageal adenocarcinoma (EAC) [28], but our optimal design framework developed here contributes unique strengths based on theory, and can be used to complement and enhance these current methods of screening design assessment. In particular, while microsimulation analyses can evaluate the benefits and harms of many hypothetical scenarios, this process can be time-intensive, prone to sensitivities in parameter choices, and ultimately only test a finite number of options [48]. In contrast, we showed that the probabilities of successful population screening can be analytically solved in certain cases and that these equations 1) are straightforward to optimize in a single computation, 2) reveal the structure of parameters and model assumptions that have the greatest implications on the sensitivity of the results obtained, and 3) predict future trends to obtain results for younger birth cohorts in a logical, data-driven way.

In the case of BE, screening has been recommended to begin in patients with multiple risk factors including increasing age (the specific threshold is 50 years of age), but the clinical evidence for these recommendations have been low to moderate in quality [31]. In order to justify the proposed recommendations, cost-effectiveness studies have used decision analytic models to quantify expected costs accrued versus life-years gained via simulations of initial screening and subsequent surveillance including endoscopic eradication therapy of detected neoplasia. While these models have each typically tested a small number of screening cohorts, we considered the full range of screening ages and found that the *optimal ages* to begin screening patients with symptomatic gastroesophageal reflux disease (GERD) were age 58 for men and 64 for women. These ages are still within cost-effective ranges found for men with GERD in previous modeling analyses that compared a few possible screening ages (including 50, 60, and 70) in sensitivity analyses [16, 49], particularly when using minimally invasive, cheaper screening modalities even in women [16]. With our framework we also reasoned that earlier screening ages are more costly, although the cost function defined serves as a simple example for how optimal control functions in cancer control programs could quantify trade-offs of futile surveillance with underdiagnoses, but extensions could also include quality-adjusted life years (QALYs).

Using our framework in a similar way, we also determined that if BE screening were recommended in the general population, the optimal ages to recommend would be age 64 for men and age 69 for women (for equal weighting given to BE onset and EAC risk in optimization). These findings for the ages of optimal BE yield are supported by prospective studies’ results that it is more likely to find BE during screening at ages closer to 60 than 50 [50], specifically in two large studies that found the same mean age at BE diagnosis of 61 in men [51], and 61 in both genders together [52]. Lastly, women found with BE were more likely to be older, between ages 61 to 70 at diagnosis, than women found without BE [53, 54]. Taken together, these studies confirm our findings that it may be justified biologically to begin screening at these later ages rather than worrying a large portion of the population starting at age 50. These proposed older ages to initialize screening in GERD and general population could be specifically tested in future cost-effectiveness analyses.

Our model describes a number of key transitions in cell biology, each defined by a specific rate parameter. Direct measurement of these parameters is usually lacking, and indeed there remains uncertainty about what key rate-limiting transitions in cancer evolution are essential. Nevertheless, our model is broadly resistant to this uncertainty as we found in the tightly estimated range for optimal screening ages in our sensitivity analyses, because the main timescales and windows of opportunity to target in our strategies are robust to perturbations in cell-scale variables to ensure that we recapitulate expected cancer incidence. First, the model’s stages were determined via rigorous likelihood ratio testing when fit to SEER cancer registry data, and its structure is further supported by whole-genome sequencing data pointing to the existence of boundary-defining mutations (or combinations of mutations) in BE that create windows for curative interventions [55]. We then must make certain assumptions - for example that the first two ‘hits’ of initiation of high grade dysplasia in BE (e.g., inactivation of each allele of *TP53*) are equally likely because we can only infer the product of the two rates with this method. Although our application does not explicitly include molecular data, we note that these parameters could be more uniquely measured using multi-region samples of clones within patient BE segments over time, especially as resources like Pre-Cancer Genome Atlases are becoming available to the public [56].

With respect to effective surveillance, we did find that molecular age of precursors (beyond patient chronological age at time of screening) was a critical variable for accurate cancer risk prediction given screening outcome. Specifically, the effect of BE dwell time (time since onset to clinical diagnosis) can be built into a model for optimizing screening and we urge the importance of finding innovative means (such as molecular clocks with patient-specific data) to measure these “hidden” events that currently cannot be clinically ascertained by traditional methods [57]. Because the framework is predicated on a biological description of the natural history of cancer, the inferences drawn are predictive and will need to be validated prospectively even though the underlying mechanistic model framework was validated using available screening data. This approach complements the development of adaptive treatment strategies (ATS) and dynamic treatment regimen (DTR) that rely on large amounts of patient-specific health records and machine learning [58]. Although these new developments are promising and are likely to yield important insights considering the wealth of clinical records for common diseases, they are difficult to extend to rare or understudied diseases and therefore are prone to biases. In contrast, models borrow strength from incidence patterns and are easily interpretable and applicable to balance risks with benefits when there is an intention to treat or intervene.

While it is difficult to justify delaying screening from a public health perspective, the problem of overdiagnosis is only becoming larger as fervor for early detection assays is growing. Our study’s predictions are reasonable considering the scarcity of epidemiological data to suggest earlier screening is ideal. We show that using a mechanism-based model of cancer could help relieve the costs of unnecessary screening and surveillance, though further validation of associated risks and sensitivities is needed. Overall such approaches are currently under-utilized in public health policies, but there is clear potential to improve guideline formation through deeper mathematical examination of data for cancers where screening can be life-saving. As “learning cancer screening programs” are being proposed to test many potential strategies in populations via randomized screening arms [17], we suggest that incorporating a model-based approach from the outset could aid in an optimal design plan (such as screening age choice in certain risk strata) in order to improve health outcomes faster. We believe that innovative teamwork between physicians, mathematical modelers, and policy-makers can improve screening design thereby reducing future cancer mortality.

## 1. Acknowledgements

This research was supported by the National Cancer Institute (www.cancer.gov) under grants U01 CA152926 (CISNET), U01 CA182940 (BG-U01), U54CA163059 (BETRNET), and U01CA199336 (CISNET), and UKRI Rutherford Fund Fellowship (KC). The authors also thankfully acknowledge the contributions of raw data on Barrett’s Esophagus prevalence from the National Endoscopic Database of the Clinical Outcomes Research Initiative (CORI).

## Supplementary Figures

**Figure S1:**
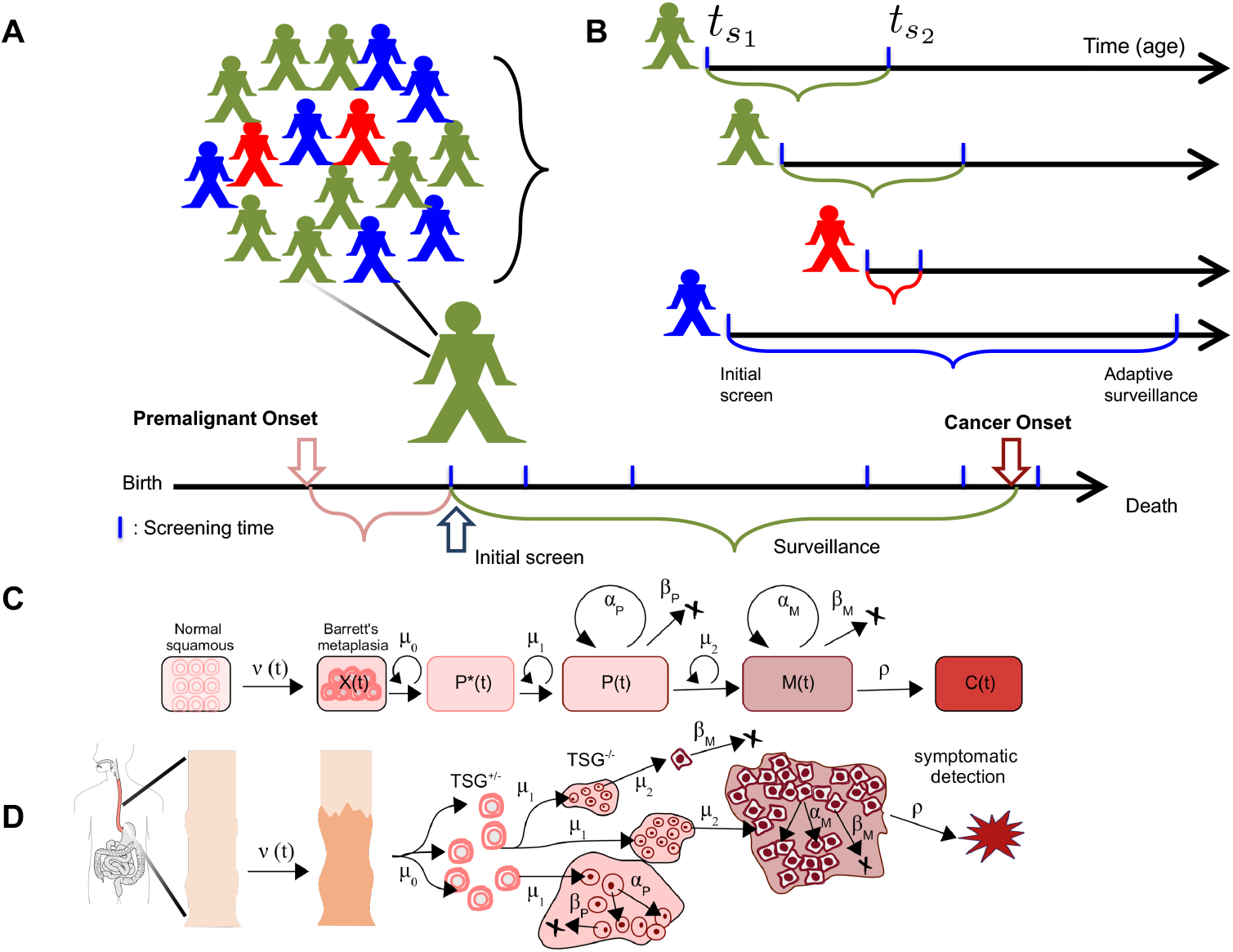
Optimizing screening and surveillance using the multistage clonal expansion model for EAC (MSCE-EAC) A) The goal of early detection and prevention is to perform an initial screen on individuals within a larger population after premalignant changes reside in cells but before incidental cancer occurs in order to intervene. B) After an initial screen at time *t*_*s*_1__, most surveillance recommendations are “one-size-fits-all” based solely on a population average (green) of a single variable-the diagnosis at initial screen (such as detection of Barrett’s esophahus (BE)). More refined adaptive recommendations for a follow-up surveillance at patient-specific time *t*_*s*_2__ can iteratively account for the heterogeneity in a screened population (i.e., other factors that contribute to a patient being low risk (blue) versus high risk (red)). C) Within the MSCE-EAC multi-type branching process framework in a single patient’s esophagus, normal squamous epithelium may transform to BE cells *X* at a random onset time with rate *ν*(*t*), followed by a ‘two-hit’ tumor initiation process with Poisson rates *μ*_0_ (creating *P*\* progeny), and *μ*_1_, which leads to the stochastic appearance of premalignant progenitor *P* cells. Premalignant cells undergo a first clonal expansion described by a birth-death-mutation process with cell division rate *α_P_*, cell death-or-differentiation rate *β_P_*, and malignant transformation rate *μ*_2_. Malignant cells *M*, in turn, may undergo a second clonal expansion by a birth-death-detection process with cell division and death rates *α_M_* and *β_M_*, respectively. Clinical detection of *M* cells occurs through a size-based detection process with parameter *ρ*. D) An illustration of MSCE-model stochastic trajectories. TSG, tumor suppressor gene; -/+, inactivation of one wild-type allele; -/-, inactivation of both wild-type alleles, e.g., loss/mutation of *TP53* in BE.

**Figure S2:**
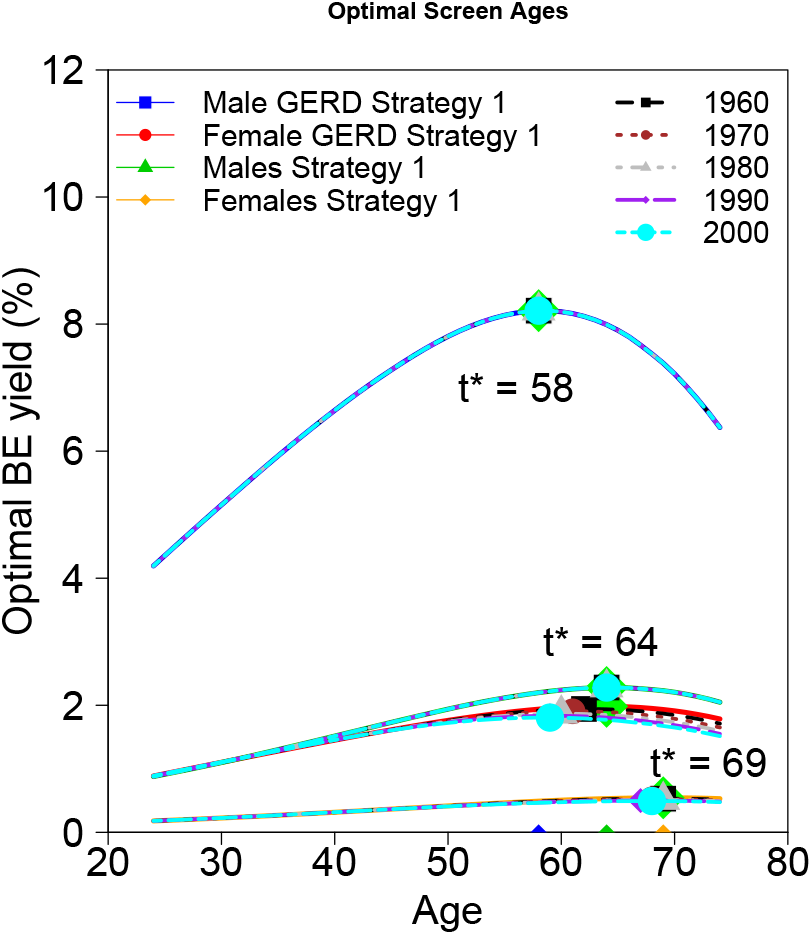
Sensitivity analysis for optimal screening ages for birth cohorts 1960 - 2000. We found that the optimal ages for men of both risk strata remain unchanged from the base case results (birth cohort 1950, Figure 4B) with increasing birth cohort. Specifically, for men with GERD we found that optimal screening age was 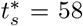 and for all men combined was 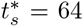 for all 5 additional birth cohorts tested. For women, we found slight differences due to the flatter optimality criterion computed: optimal initial screening ages were 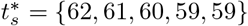 for women with GERD, and 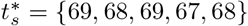 for all women combined, for birth cohorts 1960, 1970, 1980, 1990, and 2000, respectively, denoted by color.

## Supplementary Tables

**Table S1:** MSCE-EAC model biological parameters (rates per cell per year).

| Value (95% CI) | Males | Females |
| --- | --- | --- |
| $\nu_0$ | $3.65 (3.19 - 4.13) \times 10^{-4}$ | $7.48 (4.87 - 10.29) \times 10^{-5}$ |
| $\mu_0(\mu_1)$ | $7.99 (6.38 - 9.83) \times 10^{-4}$ | $7.05 (6.13 - 12.25) \times 10^{-4}$ |
| $\mu_2$ | $4.54 (3.65 - 6.47) \times 10^{-5}$ | $6.89 (3.16 - 14.28) \times 10^{-5}$ |
| $g_{P,0}^*$ | $9.91 (9.28 - 10.99) \times 10^{-2}$ | $1.23 (1.06 - 1.35) \times 10^{-1}$ |
| $g_1^*$ | $5.09 (2.75 - 5.90) \times 10^{-1}$ | $6.40 (2.16 - 8.44) \times 10^{-1}$ |
| $g_2^*$ | $5.38 (4.83 - 5.72) \times 10^{-2}$ | $2.98 (2.47 - 3.44) \times 10^{-2}$ |
| $g_3^*$ | $1912.5 (1909.1 - 1914.1)$ | $1945.3 (1923.9 - 1954.4)$ |

\*Variables denoted with $g$ are parameters of sigmoidal functions specific to a birth cohort and gender to obtain premalignant and malignant clonal proliferation rates, respectively, so that solving $g_P = \alpha_P - \beta_P - \mu_2$ and $g_M = \alpha_M - \beta_M - \rho$ obtains the parameters in Fig S1 (see [25] for explicit details).

## Supplementary Methods

### 1. Multistage clonal expansion (MSCE) model survival functions

We first introduce the notation for the following random variables of the multi-type branching process

*BE*(*t*) = Bernoulli random variable for BE conversion by time *t*
*X*(*t*) = number of BE stem cells in a tissue at time *t*
*P*\*(*t*) = number of pre-initiated cells at time *t*
*P*(*t*) = number of premalignant (initiated) cells at time *t*
*M*(*t*) = number of malignant (preclinical) cells (prior to detection) at time *t*
*C*(*t*) = number of cancer cells (after detection) at time *t*
*D*(*t*) = Bernoulli random variable for clinical detection by time *t*

Let us consider the probability generating function (pgf) Ψ for the entire process starting at *τ* = 0, ie. when an individual is born

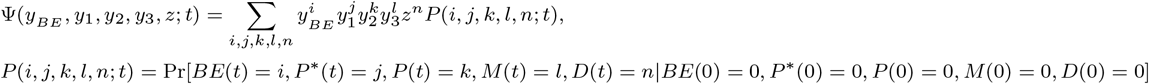

where, explicitly, *i*, *n* = {0,1} and *BE*(*t*), *D*(*t*) are the following indicator functions corresponding to BE conversion and EAC clinical detection, respectively

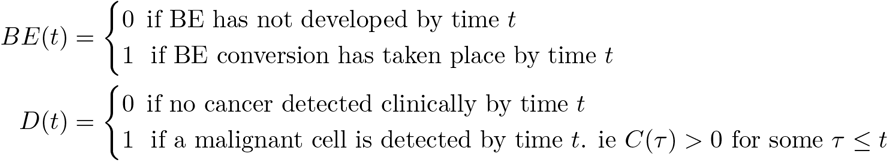

The Chapman-Kolmogorov equations governing the transition probabilities for this multistage process include contributions from the initial Armitage-Doll type transition to BE, the two Poisson transitions to initiation, and the two birth-death-migration processes, all of which have been derived previously [13, 59, 60]. We begin with a method for solving for these generating functions using the Kolmogorov backward equations. Beginning with an active BE segment (*BE*), a single pre-initiated (*P*\*), premalignant (*P*), or malignant (*M*) cell at time *τ* only, we define the following generating functions Φ_*BE*_,Φ_*P*_, Φ_*P*_, or Φ_*M*_, respectively,

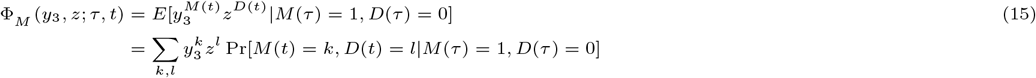

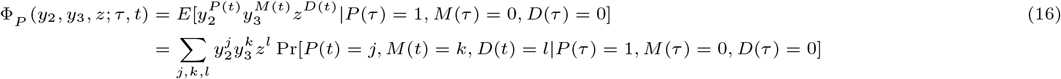

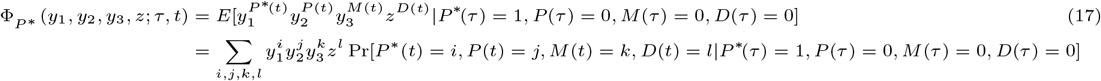

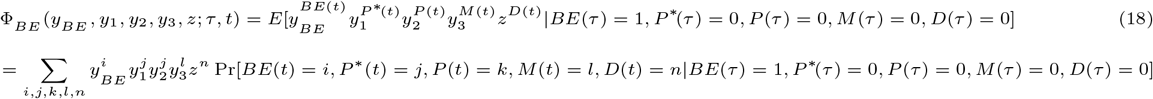

The generating functions satisfy the following Kolmogorov backward equations

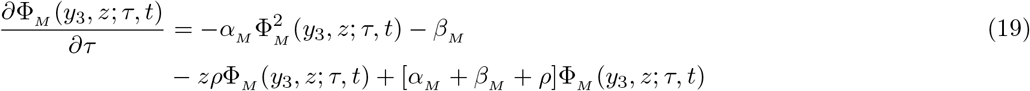

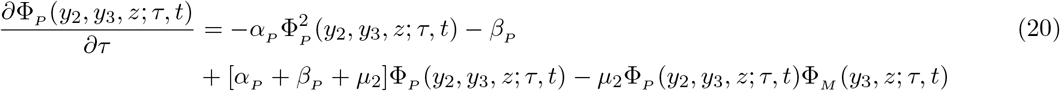

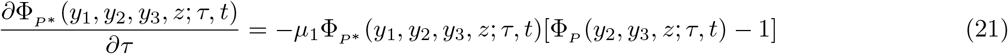

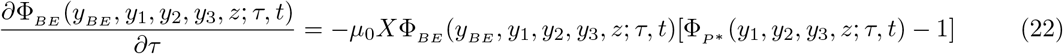

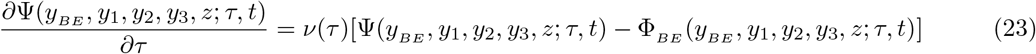

To connect the cellular level description to the population level, we first solve for the overall survival function (for EAC cancer detection), starting at time 0, which in our notation is

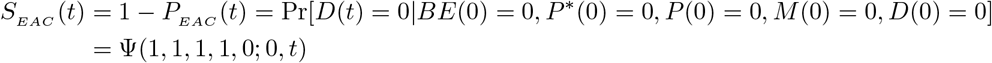

where *P_EAC_*(*t*) is the probability of a cancer detection at time *t*,

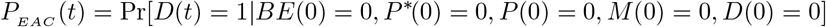

We will here denote Φ_*M*_(1,0; *τ, t*) ≡ Φ_*M*_(*τ*, *t*), Φ_*P*_(1,1, 0; *τ*, *t*) ≡ Φ_*P*_ (*τ*, *t*), Φ_*P*\*_(1,1,1,0; *τ*, *t*) ≡ Φ_*P*\*_(*τ*, *t*), Φ_*BE*_(1,1,1,1, 0; *τ*, *t*) ≡ Φ_*BE*_(*τ*, *t*), and Ψ(1,1,1,1, 0; *τ*, *t*) ≡ Ψ(*τ*, *t*). A dot designates a first derivative with respect to *t*. The hazard function, i.e., the rate at which cancer is detected in individuals who have not been diagnosed before, is given by

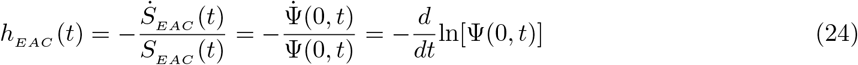

For fixed *t*, this boundary value system of coupled PDEs can be converted into an initial value problem (IVP) with the change of variables *u* = *t* − *τ*, where *u* is the “running” time. This redefinition and equations hereafter follow the method used by Crump et al. [61]. Define the following variables for the new IVP: 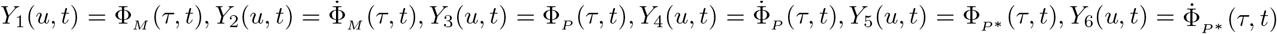,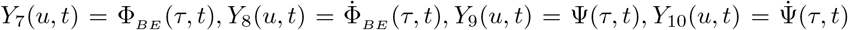 with corresponding initial conditions *Y*_1_(0,*t*) = *Y*_3_(0,*t*) = *Y*_5_(0,*t*) = *Y*_7_(0,*t*) = *Y*_9_(0,*t*) = 1, *Y*_4_(0,*t*) = *Y*_6_(0,*t*) = *Y*_8_(0,*t*) = *Y*_10_(0,*t*) = 0, and *Y*_2_(0,*t*) = −*ρ*. Then the equations to solve for our IVP are the following

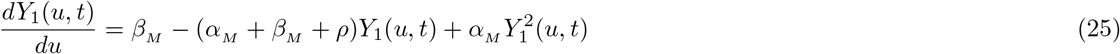

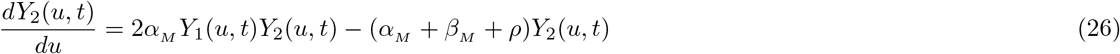

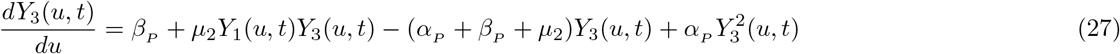

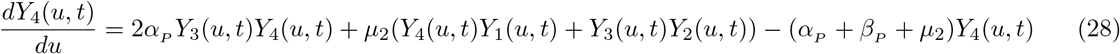

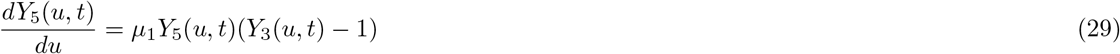

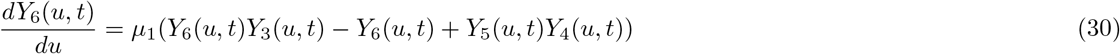

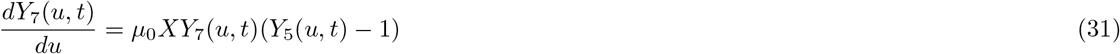

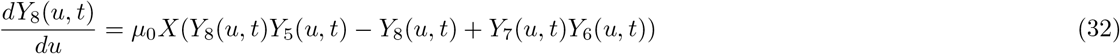

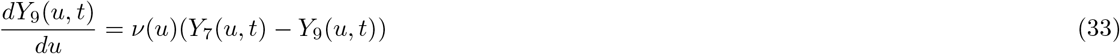

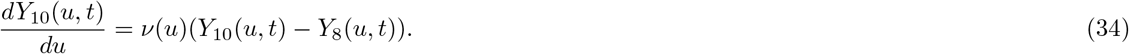

These 10 coupled ODEs can be solved numerically to obtain the desired survival functions,

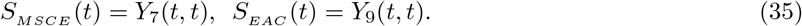

In the Results, we assigned binary detection variables the value 1 if a clone born at time s is non-extinct at time *t_s_* (size of clone is ≥ 1) and 0 otherwise. In the case of EAC, this “perfect sensitivity” definition will be useful as progressively better high-resolution imaging technologies and minimally invasive sampling devices are tested in trials in BE [16, 30] and used in future clinical practice. For detectability based on this non-extinction criterion at time of screening, we adopt the probability of detection with a binary outcome variable for malignant clones with an unconditional birth-death-mutation size distribution (see Jeon et al. [18]). The forms of these are given by,

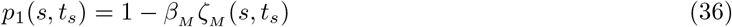

where

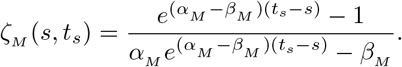

As *t_s_* → ∞, *p*_1_(*s,t_s_*) → 1 − *β_M_*/*α_M_*. We obtain the analogous definition for *p*_2_(*s,t_s_*) with *P* parameters for the detection probability of premalignant clones -

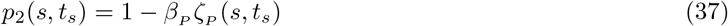

where

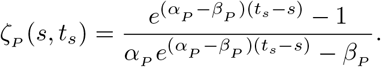

As *t_s_* → ∞, *p*_2_(*s,t_s_*) → 1 − *β_P_*/*α_P_*, which is the asymptotic probability of non-extinction.

### 2. Initial screen strategy 2

We aim to derive the optimal screening age 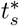 (Eq. (8) in Main Text)

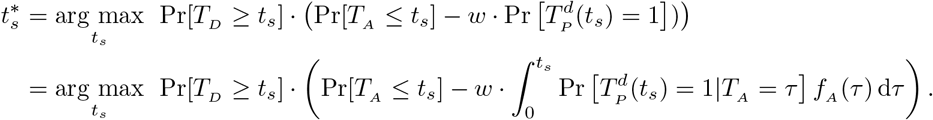

Applying useful theory in multi-type branching processes for multistage models, we utilize the filtered Poisson process (FPP) approach [62, 63]) to analytically solve for 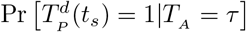. Let *N*(*σ, t_s_*) be the number of initiated *P* cells from a *P*\* cell born at time *σ* that result in a detectable HGD clone at time *t_s_*

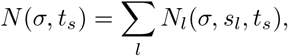

where *N_l_*(*σ, t_s_, s_l_*) = 1 if initiation at time *s_l_* produces a detectable clone at time *t_s_*, and *s_l_* ≤ *t_s_*, with probability 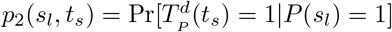. Then *N*(*σ,t_s_*) is a FPP such that

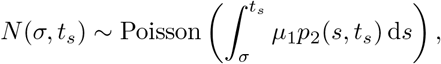

where *p*_2_(*s_l_,t_s_*) is defined in Eq. (37). Next let *M*(*τ, t_s_*) be the number of preinitiated cells after BE onset at time *τ* that result in a detectable P clone at time *t_s_*. Then we have

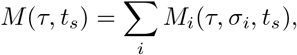

where *M_i_*(*τ,σ_i_,t_s_*) is binary and is equal to 1 if the preinitiated cell originting at time *σ_i_* after BE onset at time *τ* results in a detectable P clone at time *t_s_*, with *τ* ≤ *σ_i_* ≤ *t_s_*. Note that *M_i_*(*τ, σ_i_, t_s_*) = 1 is equivalent to *N*(*σ_i_,t_s_*) ≥ 1 which has the probability 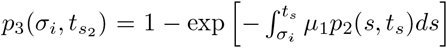 with *p*_2_(*s,t_s_*) given by Eq. (37). Then *M*(*τ,t_s_*) is a FPP such that

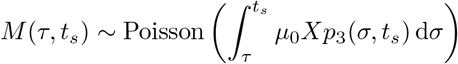

Thus

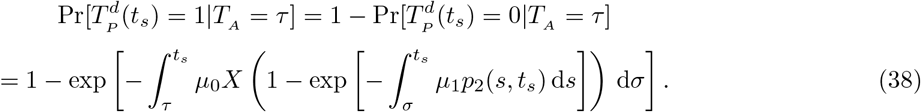

Thus together with 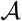 onset distribution in Eq. (5) and mortality risk, we have the analytical solution for optimal screening age in Strategy 2, given in Eq. (8).

### 3. Initial screen strategy 3

Here we derive the analytical solution for the optimal screening time for Strategy 3, given some weighting *w* to avoid risk of co-existing malignant clones (Eq. (9) in Main Text),

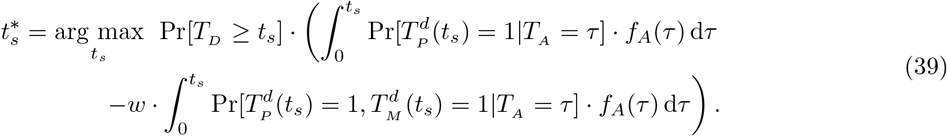

The first integral in this difference was derived in Strategy 2 (see Eq. (38)). We solve for the probability in the integrand of the second integral with detectability given in Eq. (36) and Eq. (37) for malignant clones and HGD clones, respectively. Thus we seek an explicit formula for the integrand,

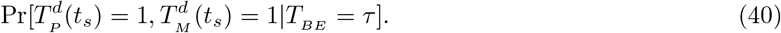

To derive the above probability, we must define *C*_1_, *C*_2_, *C*_3_, mutually exclusive scenarios of counts of initiations of different types. Scenario *C*_1_ corresponds to the scenario that the *P* clone that is non-extinct at time *t_s_* is the ancestor of the non-extinct *M* clone at time *t_s_*. This is most commonly the case in the stochastic realizations since the bigger *P* clones that remain non-extinct at time of screening have the highest probability to have had *μ*_2_ events (malignant transformation) occur since their initiation. Scenario *C*_2_ accounts for a non-extinct *P* clone at time *t_s_* that, if it created any malignant progeny before time of screening, those malignancies went extinct. Scenario *C*_3_ accounts for a non-extinct *M* clone at time *t_s_* whose ancestral *P* clone went extinct before time t_s_. Thus the probability we aim to derive accounts for either event *C*_1_ or *C*_2_ and together. For each of the three scenarios, we will use filtered Poisson processes techniques and abuse notation slightly by redefining FPP random variables *N*_1_ and *N*_2_ differently but analogously for each scenario, to avoid verboseness. We start with deriving the probability that event *C*_1_ occurs (at least once) with *u* > *s*,

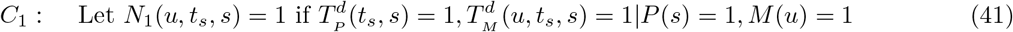

with probability 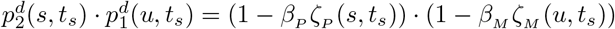. Then we have that

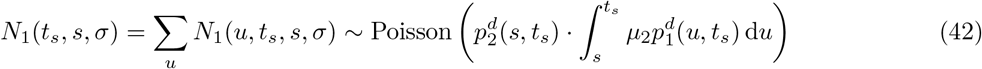

and

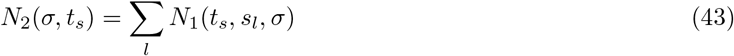

so that *N*_2_(*σ,t_s_*) is a FPP and the probability generating function (PGF) of *N*_2_(*σ,t_s_*) is

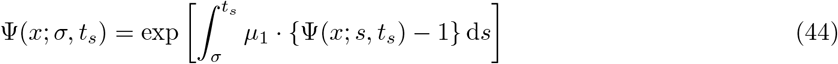

where Ψ(*x*; *s*, *t_s_*) is the PGF of *N*_1_(*t_s_, s, σ*) = *e*^−λ^(1−*x*), where 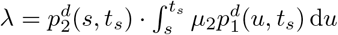. Therefore

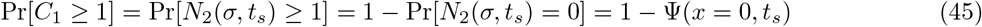

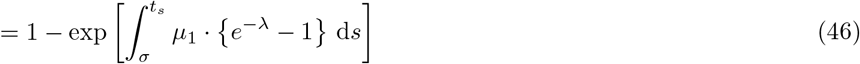

Similarly,

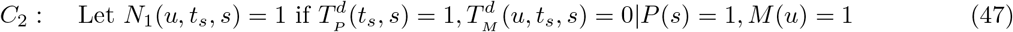

with probability 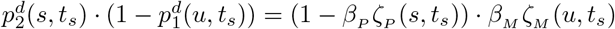. Then we have that

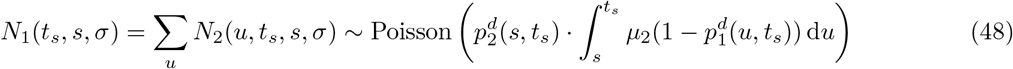

and

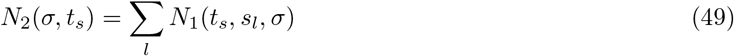

so that *N*_2_(*σ, t_s_*) is a FPP and the PGF of *N*_2_(*σ, t_s_*) is

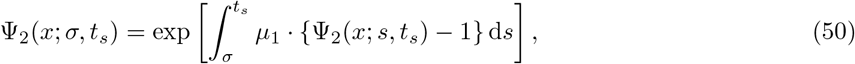

where Ψ_2_(*x*; *s, t_s_*) is the PGF of *N*_1_(*t_s_,s,σ*) = *e*^−λ_2_(1−*x*)^, where 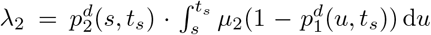. Therefore

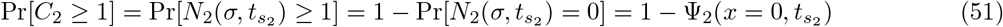

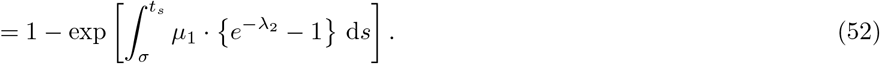

Lastly, we define

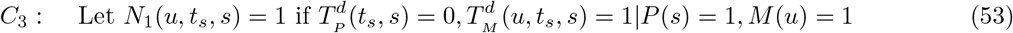

with probability 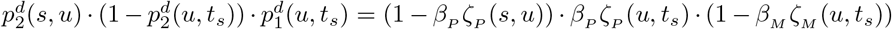 Then we have that

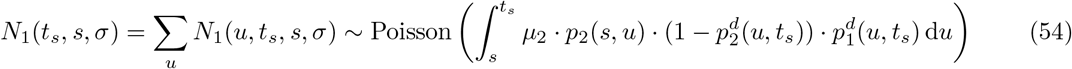

and

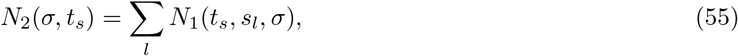

so that *N*_2_(*σ, t_s_*) is a FPP and the PGF of *N*_2_(*σ, t_s_*) is

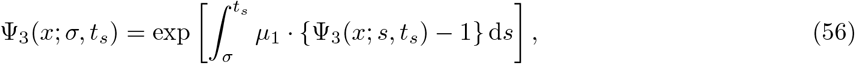

where Ψ_3_(*x; s, t_s_*) is the PGF of *N*_1_(*t_s_, s, σ*) = *e*^−λ_3_(1−*x*)^, where 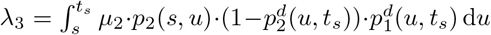. Therefore

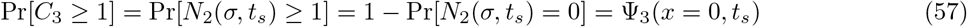

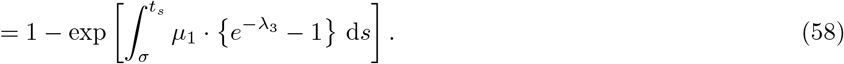

Next, let *M*(*τ, t_s_*) be the number of preinitiated cells after BE onset at time *τ* that result in *C*_1_ ∪ (*C*_2_ ∩ *C*_3_)

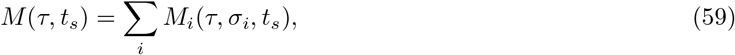

where *M_i_*(*τ, σ_i_, t_s_*) = 1 ⇔ *C*_1_ ≥ 1 ∪ (*C*_2_ ≥ 1, *C*_3_ ≥ 1). Due to the mutual exclusivity of these events, we have that *M_i_*(*τ, σ_i_, t_s_*) = 1 with probability

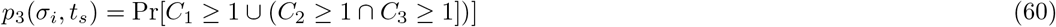

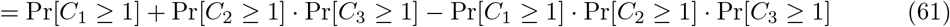

Then *M* (*τ, t_s_*) is a FPP such that

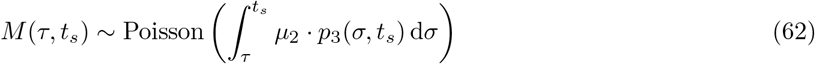

Finally, we solve for our integrand of interest from Eq. (40)

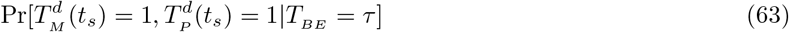

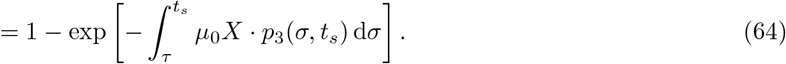

With this, we have analytically solved for all components of the optimality criterion for Strategy 3 from Eq. (39).

### 4. Adaptive surveillance and associated-risk

Continuing from the Main Text we derive the following for optimal time for surveillance given Outcome 1 (negative for precursor),

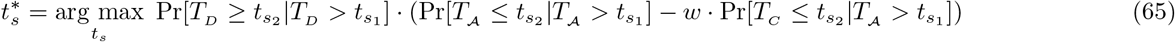

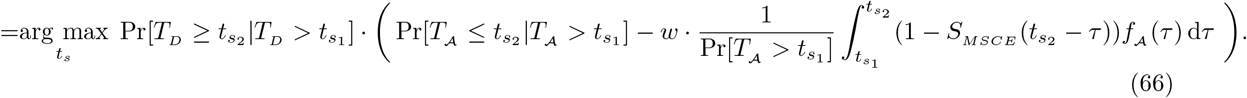

Then for Outcome 2 (positive for precursor) we defined a risk associated with a surveillance timing as,

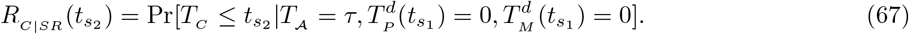

Here we derive this risk associated with next surveillance time given a certain screening result, *R_C|SR_*(*t*_*s*_2__), as general lifetime risk without other cause mortality for use in optimization/cost-effectiveness analysis. Once again, for the scenario with a perfectly sensitive screen that can detect single cells, we may derive this probability using a filtered Poisson approach analogous to the approach used in Strategy 2 for a single screen. We redefine (for sake of retaining simpler notation) *N*(*σ, t*_*s*_2__) to be the number of initiated *P* cells from a *P*\* cell born at time *σ* that result in a clinically detected EAC clone at time *t*_*s*_2__

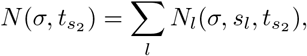

where *N_l_*(*σ, s_l_,t*_*s*_2__) = 1 if initiation at time *s_l_* produces EAC at time *t*_*s*_2__, and *t*_*s*_1__ < *s_l_* < *t*_*s*_2__, with probability *p_C_* (*s_l_*, *t*_*s*_2__) = Pr[*T_C_* < *t*_*s*_2__ |*P*(*s_l_*) = 1]. Then *N* (*σ,t*_*s*_2__) is a FPP such that

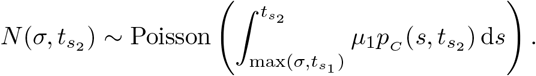

Clinical EAC cumulative distribution, *p_C_* (*s_l_,t*_*s*_2__), can be solved via the Kolmogorov equations of the MSCE-EAC branching process from the PDE in Eq. (20). Specifically we have

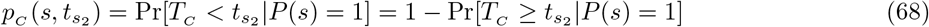

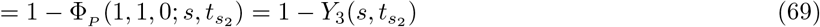

where Φ_*p*_(1,1,0; *s,t*_*s*_2__) is the EAC survival from a single *P* cell (see Eq. (16)), *Y*_3_(*u,t*_*s*_2__) is the associated ODE (see Supplementary Methods 1) and *u* = *t*_*s*_2__ – *s*. Next let *M*(*τ, t*_*s*_2__) be the number of preinitiated cells after BE onset at time *τ* that result in detectable EAC at time *t*_*s*_2__. Then we have

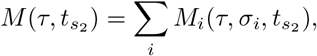

where *M_i_*(*τ, σ_i_,t*_*s*_2__) is, as before, binary and is equal to 1 if the preinitiated cell originting at time *σ_i_* after BE onset at time *τ* results in a detectable EAC at time *t*_*s*_2__, with *τ* < *t*_*s*_1__ < *t*_*s*_2__. Note that *M_i_*(*τ,σ_i_,t*_*s*_2__) = 1 is equivalent to *N*(*σ_i_,t*_*s*2_) ≥ 1 which has the probability 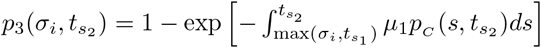. Then *M*(*τ, t*_*s*_2__) is a FPP such that

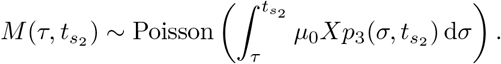

Thus

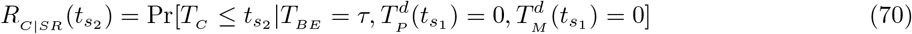

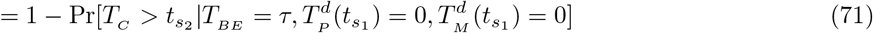

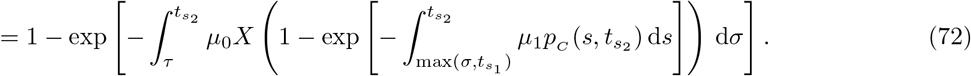

### 5. Adaptive screen outcome 2, strategy 1

As a final example of adaptive surveillance, an important clinical question to consider is, “Should we offer a one-time surveillance and when would that optimal time be?” Assuming we have an estimate *τ* of a patient’s precursor onset time, we solve for optimal next-surveillance age 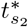,

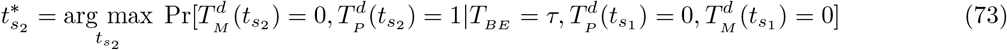

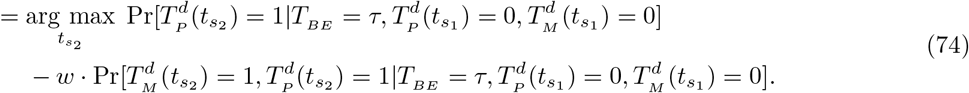

Again, we can increase or decrease *w* depending on how much weight we want to place on keeping the risk of developing malignancy under a certain value, say *ϵ*. For ease of notation in this strategy, we do not include the other cause mortality term Pr[*T_D_* ≥ *t*_*s*_2__ |*T_D_* > *t*_*s*_1__] that can be multiplied to the entire expression of optimality criterion provided in Eq. (74) as we have done in previous strategies.

Let us first derive a formula for the first term of this difference based on the MSCE-EAC model construction using again a filtered Poisson process approach,

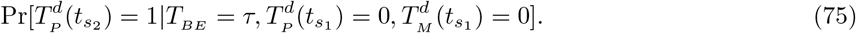

Let *N*(*σ, t*_*s*_1__, *t*_*s*_2__) be the number of initiated cells from a *P*\* cell born at time *σ* that result in a detectable clone at time *t*_*s*_2__, not born before *t*_*s*_1__

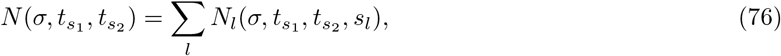

where *N_l_*(*σ,t*_*s*_1__,*t*_*s*_2__, *s_l_*) = 1 if initiation at time *s_l_* produces a detectable clone at time *t*_*s*_2__, *s_l_* > *t*_*s*_1__ with probability 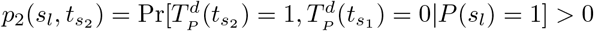 if and only if *s_l_* > *t*_*s*_1__.

Then *N*(*σ, t*_*s*_1__, *t*_*s*_2__) is a FPP such that

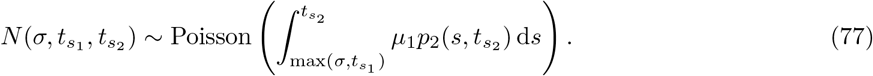

Assuming detectability of *P* cells based on non-extinction at time of screening, we have the following probability (see Eq. (37)),

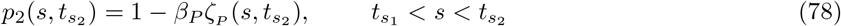

where

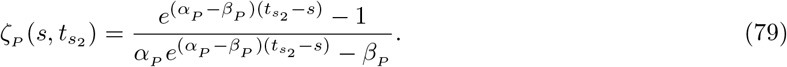

The asymptotic solution for this probability is *p*_2_ (*s, t*_*s*_2__) = 1 – *β_P_*/*α_P_*. We will assume the same for malignant clones, with detection probability provided in Eq. (36).

Next let *M*(*τ, t*_*s*_2__) be the number of preinitiated cells after BE onset at time *τ* that result in a detectable P clone at time *t*_*s*_2__. Then we have that

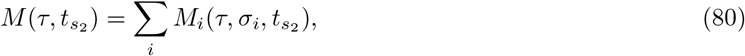

where *M_i_*(*τ, σ_i_,t*_*s*_2__) is, as before, binary and is equal to 1 if the preinitiated cell originting at time *σ_i_* after BE onset at time *τ* results in a detectable EAC at time *t*_*s*_2__, with *τ* < *t*_*s*_1__ < *t*_*s*_2__. Note that *M_i_*(*τ,σ_i_,t*_*s*_2__) = 1 is equivalent to *N*(*σ_i_,t*_*s*_2__) ≥ 1 which has the probability 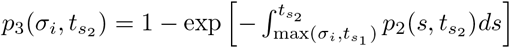.

Then *M*(*τ, t*_*s*_2__) is a FPP such that

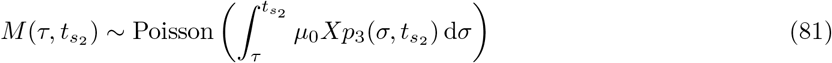

Thus

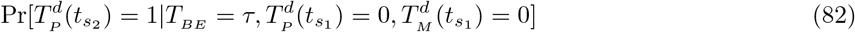

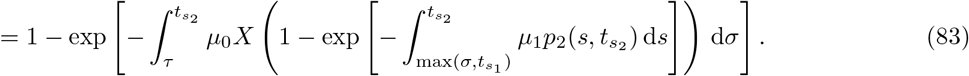

For the second term in Eq. (74), we will now derive

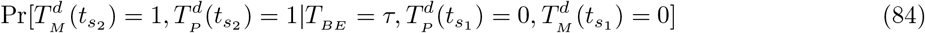

We define similar *C*_1_,*C*_2_,*C*_3_ mutually exclusive scenarios of counts of initiations of different types as was utilized in initial screen Strategy 3. For each of the three scenarios, we will use filtered Poisson processes techniques and abuse notation slightly by redefining FPP random variables *N*_1_ and *N*_2_ variables differently but analogously for each scenario, to avoid verboseness. We start with deriving the probability that event *C*_1_ occurs (at least once),

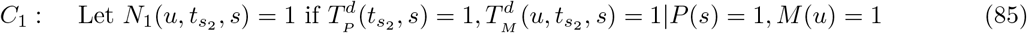

with probability 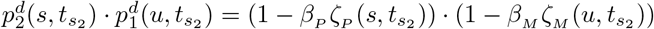. Then we have that

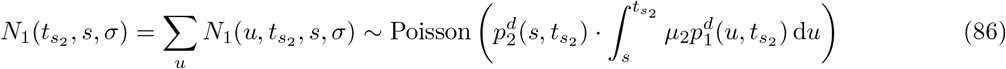

and

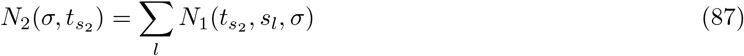

so that *N*_2_(*σ, t*_*s*_2__) is a FPP and the PGF of *N*_2_ (*σ, t*_*s*_2__) is

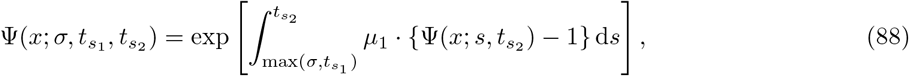

where Ψ(*x; s,t*_*s*_2__) is the PGF of *N*_1_(*t*_*s*_2__; *s; σ*) = *e*^−*λ*(1–*x*)^ where 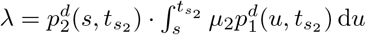 Therefore

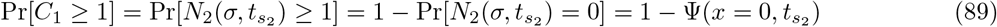

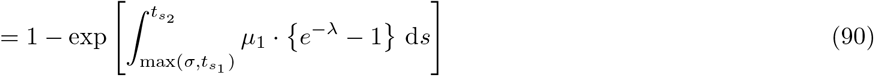

Similarly,

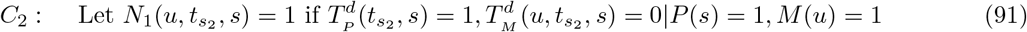

with probability 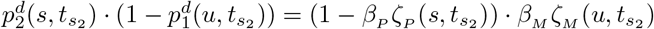. Then we have that

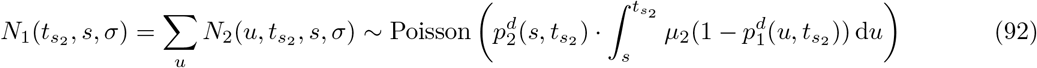

and

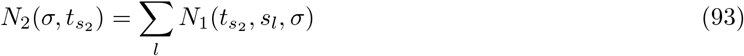

so that *N*_2_(*σ,t*_*s*_2__) is a FPP and the PGF of *N*_2_(*σ,t*_*s*_2__) is

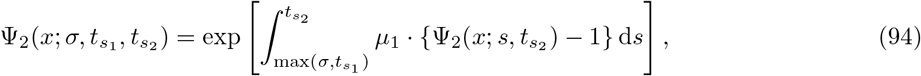

where Ψ^2^(*x*; *s,t*_*s*_2__) is the PGF of *N*_1_(*t*_*s*_2__, *s,σ*) = *e*^−λ_2_(1 – *x*)^ where 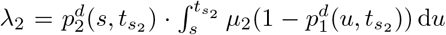. Therefore

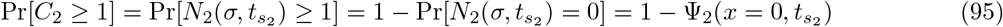

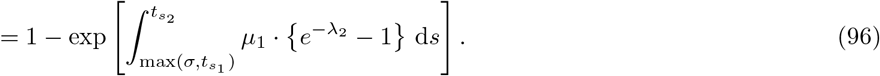

Lastly, we define

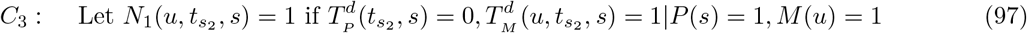

with probability 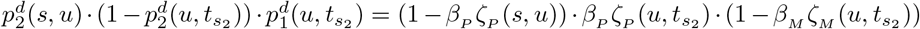. Then we have that

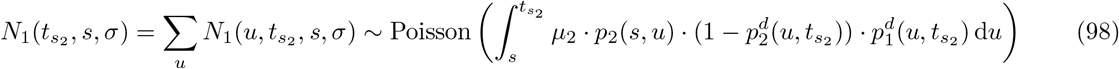

and

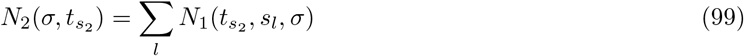

so that *N*_2_(*σ, t*_*s*_2__) is a FPP and the PGF of *N*_2_(*σ, t*_*s*_2__) is

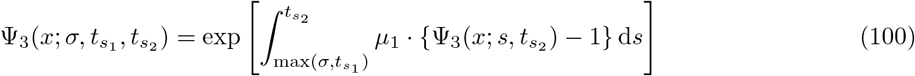

where Ψ_3_(*x; s,t*_*s*_2__) is the PGF of *N*_1_(*t*_*s*_2__,*s,σ*) = *e*^-λ_3_(1–*x*)^ where 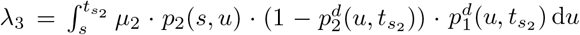. Therefore

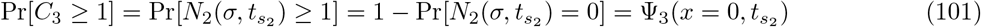

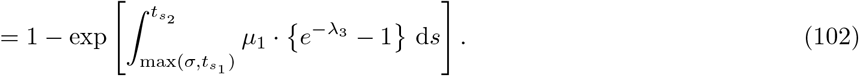

Next, let *M*(*τ, t*_*s*_2__) be the number of preinitiated cells after BE onset at time τ that result in *C*_1_ ∪ (*C*_2_ ∩ *C*_3_)

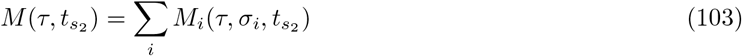

where 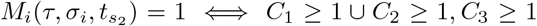. Due to the mutual exclusivity of these events, we have that *M_i_*(*τ, σ_i_,t*_*s*_2__) = 1 with probability

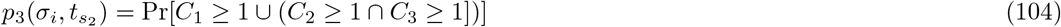

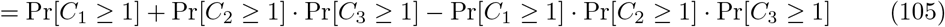

Then *M*(*τ, t*_*s*_2__) is a FPP such that

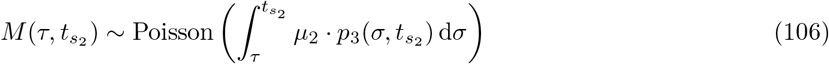

With these, we have analytically solved for our expression of interest from Eq. (84),

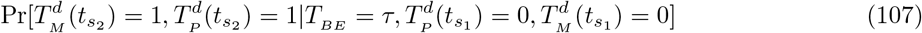

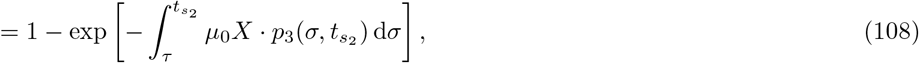

and thus have both components to compute the optimality criterion for Outcome 2: Strategy 1 from Eq. (74).

